# A longitudinal resource for population neuroscience of school-age children and adolescents in China

**DOI:** 10.1101/2023.04.07.535977

**Authors:** Xue-Ru Fan, Yin-Shan Wang, Da Chang, Ning Yang, Meng-Jie Rong, Zhe Zhang, Ye He, Xiaohui Hou, Quan Zhou, Zhu-Qing Gong, Li-Zhi Cao, Hao-Ming Dong, Jing-Jing Nie, Li-Zhen Chen, Qing Zhang, Jia-Xin Zhang, Hui-Jie Li, Min Bao, Antao Chen, Jing Chen, Xu Chen, Jinfeng Ding, Xue Dong, Yi Du, Chen Feng, Tingyong Feng, Xiaolan Fu, Li-Kun Ge, Bao Hong, Xiaomeng Hu, Wenjun Huang, Chao Jiang, Li Li, Qi Li, Su Li, Xun Liu, Fan Mo, Jiang Qiu, Xue-Quan Su, Gao-Xia Wei, Yiyang Wu, Haishuo Xia, Chao-Gan Yan, Zhi-Xiong Yan, Xiaohong Yang, Wenfang Zhang, Ke Zhao, Liqi Zhu, Lifespan Brain Chart Consortium (LBCC), Chinese Color Nest Consortium (CCNP), Xi-Nian Zuo

**Author notes:** These authors contributed equally to this work as first authors (Xue-Ru Fan, Yin-Shan Wang, Da Chang). LBCC is an international consortium and has aggregated 123,984 MRI scans, across more than 100 primary studies, from 101,457 human participants between 115 days post-conception to 100 years of age, and built brain charts to identify previously unreported neurodevelopmental milestones. More information are available at https://github.com/brainchart/lifespan. CCNP is a long-term effort (2013-2032) to build the lifespan brain-mind development cohort in China, and more consortium information are available at http://deepneuro.bnu.edu.cn/?p=163. Corresponding author(s): Xi-Nian Zuo (Website: https://zuoxinian.github.io;,; Twitter: zuoxinian).

## Abstract

During the past decade, cognitive neuroscience has been calling for population diversity to address the challenge of validity and generalizability, ushering in a new era of population neuroscience. The developing Chinese Color Nest Project (devCCNP, 2013-2022), a ten-year pilot stage of the lifespan CCNP (2013-2032), is an ongoing project focusing on brain-mind development. The project aims to create and share a large-scale, longitudinal and multimodal dataset of typically developing children and adolescents (ages 6.0–17.9 at enrolment) in the Chinese population. The devCCNP houses not only phenotypes measured by demographic, biophysical, psychological and behavioural, cognitive, affective, and ocular-tracking assessments but also neurotypes measured with magnetic resonance imaging (MRI) of brain morphometry, resting-state function, naturalistic viewing function and diffusion structure. This Data Descriptor introduces the first data release of devCCNP including a total of 864 visits from 479 participants. Herein, we provided details of the experimental design, sampling strategies, and technical validation of the devCCNP resource. We demonstrate and discuss the potential of a multicohort longitudinal design to depict normative brain growth curves from the perspective of developmental population neuroscience. The devCCNP resource is shared as part of the “Chinese Data-sharing Warehouse for *In-vivo* Imaging Brain” in the *Chinese Color Nest Project (CCNP) – Lifespan Brain-Mind Development Data Community* (https://www.scidb.cn/en/c/ccnp) at the Science Data Bank.

**Design Types:**

- Accelerated longitudinal design
- Brain-mind development
- Population imaging
- Brain chart
- Repeated measure

**Measurements:**

- Psychological behaviours
- Biophysical and physical measures
- Intelligence quotient measure
- Neuroimaging

**Sample Characteristic - Organism:**

- Homo sapiens

**Sample Characteristic - Environment:**

- School- and community-based sample

**Sample Characteristic - Location:**

- Chongqing and Beijing, China

**Duration:**

- 10 years (2013-2022)

## Background & Summary

To explore the relationship between human behaviour and the brain, especially with respect to individual differences and precision medicine, large-scale neuroimaging data collection is necessary. In 2008, thirty-five laboratories from 10 countries including China, launched the 1000 Functional Connectomes Project (FCP)^1^. This global project shared MRI data from 1,414 worldwide participants’ neuroimaging data through the Network Information Technology Resources Collaboratory (NITRC) in the United States. As a milestone in open science for human brain function, the project demonstrated the association of individual differences in functional connectivity with demographic phenotypes (age and sex)^1^. Since then, population-based prospective efforts have been implemented by worldwide brain initiatives, such as Human Connectome Project (HCP)^2^, US BRAIN Initiative (Brain Research through Advancing Innovative Neurotechnologies Initiative, or BRAIN)^3^, Brain Mapping by Integrated Neurotechnologies for Disease Studies (Brain/MINDS) in Japan^4^, UK (United Kingdom) Biobank^5^, BRAIN Canada^6^, and Adolescent Brain Cognitive Development (ABCD) study^7^. This has introduced big data into cognitive neuroscience with population imaging, namely population neuroscience^8,9^, to increase population diversity or sample representativeness for improvements in generalizability, a significant challenge faced by current cognitive neuroscience research^10,11^.

The Chinese Color Nest Project (CCNP, 2013-2032)^12^ is an early representative effort, likely the first in China, investigating brain growth during the transition period from childhood to adolescence. CCNP has built and accumulated rich and valuable experiences as a pilot study to accelerate the pace of initiating related brain-mind development cohort studies in the China Brain Project^13,14^. CCNP is devoted to collecting nationwide data on brain structure and function across different stages of human lifespan development (6–85 years old). The long-term goal of this work is to create neurobiologically sound developmental curves for the brain to characterize phenomenological changes associated with the onset of various forms of mental health and learning disorders, as well as to predict the developmental status (i.e., age-expected values) of an individual brain’s structure or function. The developmental component of CCNP (devCCNP), also known as "Growing Up in China"^15^, has established follow-up cohorts in Chongqing and Beijing, China. With the collection of longitudinal brain images and psychobehavioural samples from school-age children and adolescents (6–18 years) in multiple cohorts, devCCNP has constructed a full set of school-age brain templates, morphological growth curves^16^ and functional connectivity gradients^17^ for the Chinese Han population as well as related (although preliminary) differences in brain development between Chinese and American school-age children^16^. The project has contributed to charting human brain development across the lifespan (0–100 years) in an international teamwork led by the Lifespan Brain Chart Consortium^18^.

To expand available resources for investigating population diversity^19^ while recognizing and addressing the issues of sampling bias, and inclusion barriers within developmental population neuroscience^20^, we describe and share the brain-mind datasets of devCCNP here. We offer a comprehensive outline of the devCCNP protocol, along with recommendations to ensure that devCCNP can be scaled up to facilitate access to more diverse populations in the future. We provide all the anonymized raw data adhering to Brain Imaging Data Structure (BIDS) standards^21^. In summary, this dataset comprises ample tasks addressing neurodevelopmental milestones of both primary and higher-order cognitive functions. The dataset holds the potential to deepen our understanding of brain development in various dimensions, and augments assessments of cultural diversity among the existing datasets using accelerated longitudinal designs (ALD) (see Table 1 from the cohort profile on CCNP^12^ for a nonexhaustive list of normative developmental samples obtained by ALD). In addition, we hope that the devCCNP will provide a resource to explore potential regional differences due to multisite sampling, and their impacts on brain development.

**Table 1.**
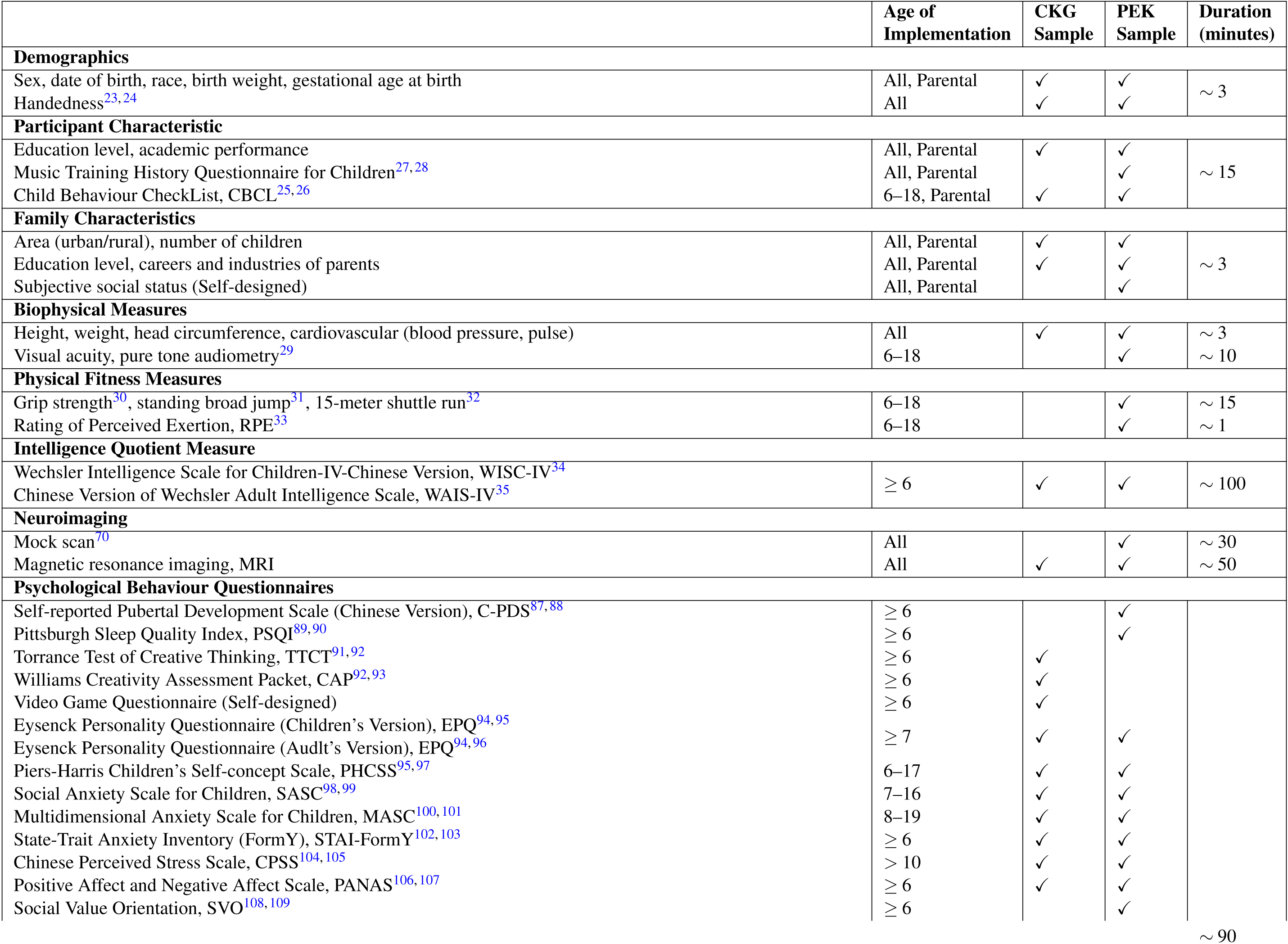
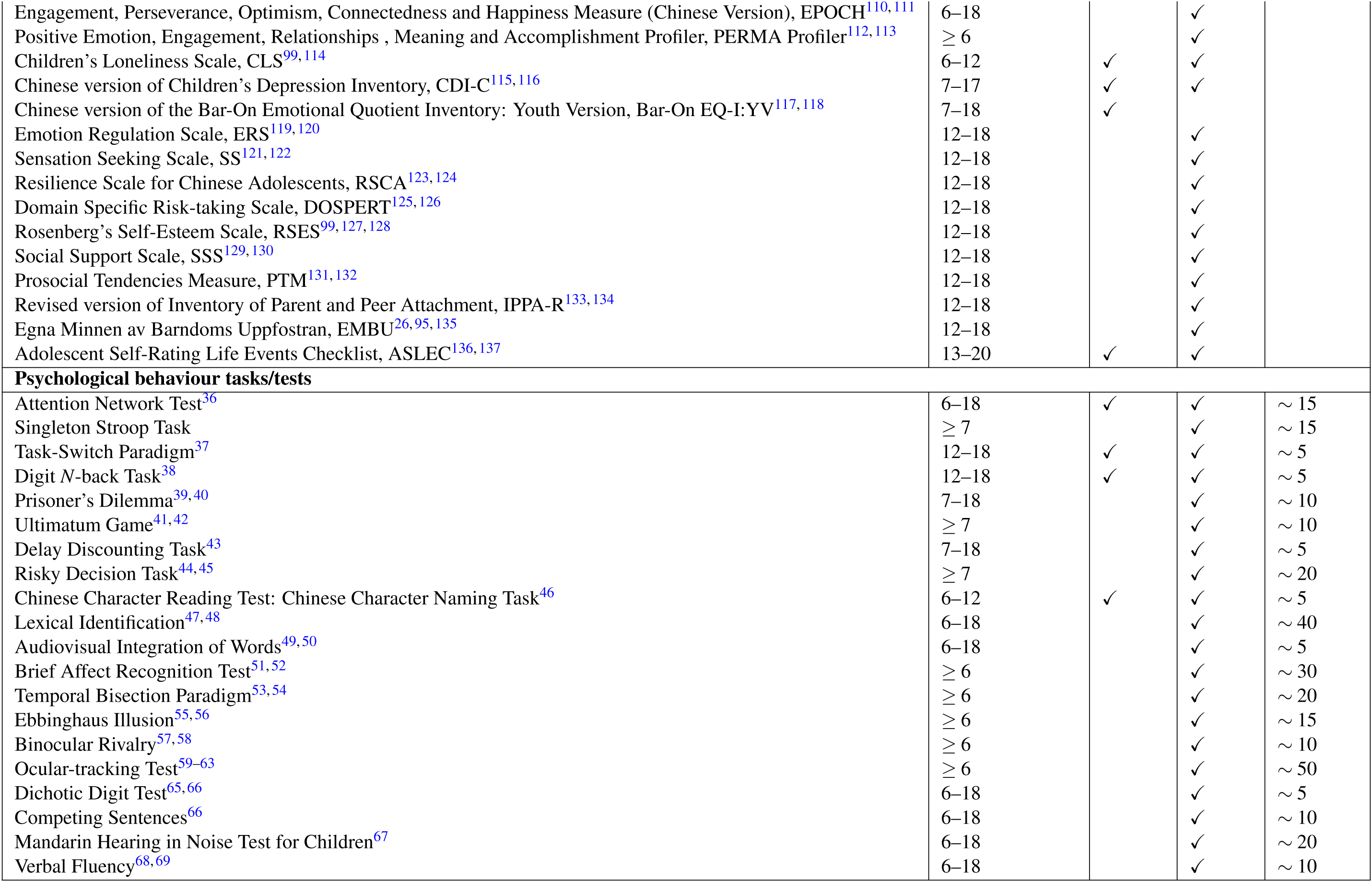
Complete protocol.

## Methods

### Overall Design

The pilot stage of devCCNP aimed to establish an ALD cohort. The cohort consisted of 480 participants with typical development, who were evenly divided into age-specific groups. Each age cohort contain 20 boys and 20 girls (Figure. 1a). We conducted data collection in two regions in China with distinct geographic and socioeconomic profiles, the Beibei District of Chongqing (devCCNP-CKG Sample) and the Chaoyang District of Beijing(devCCNP-PEK Sample), to capture a more representative sample of the Chinese population and its diverse characteristics. The devCCNP-CKG Sample was collected from March 2013 to January 2017, and the devCCNP-PEK Sample was collected from September 2017 to December 2022. Participants underwent assessment three times in total, referred to as three "waves" of visits. To account for season effects, there was a 15-month time gap between each wave (Figure. 1b). A repeated protocol was applied, which was adjusted based on the participants’ age. The total time for each assessment was approximately 10–12 hours, including preparation time and short breaks. The time duration of each visit is listed in Table 1.

**Figure 1.**
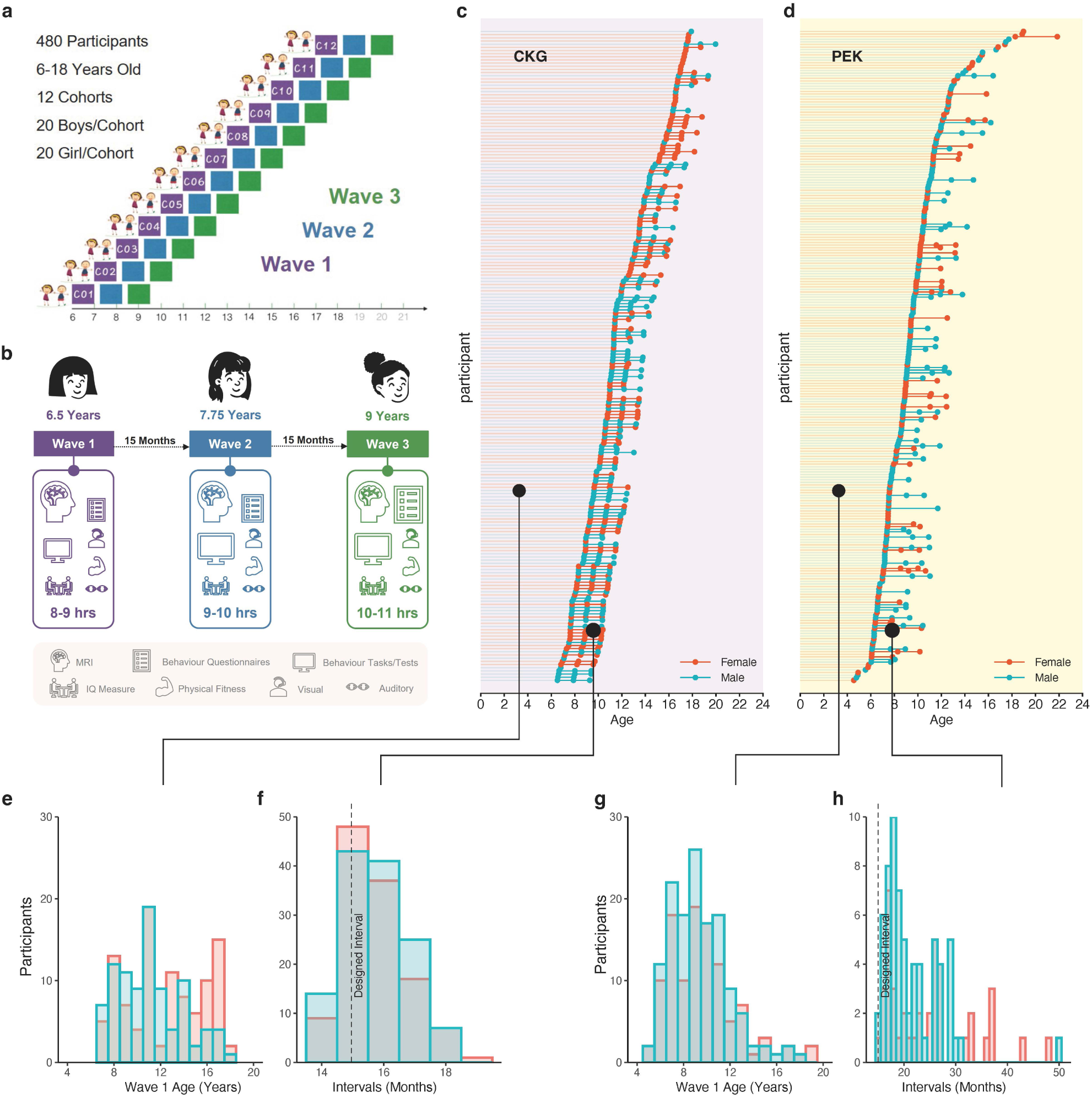
Experimental design and sample composition. (a) The Accelerated Longitudinal Design (ALD) of devCCNP has 3 repeated measuring waves: Wave 1 (baseline, purple), Wave 2 (follow-up 1, blue), and Wave 3 (follow-up 2, green). The age range of participant enrolment was 6–18 years. The 480 participants were divided into 12 age cohorts, with 20 boys and 20 girls in each. The interval between each successive waves was designed to be 15 months. (b) An example of a participant’s protocol who enrolled at 6.5 years. Measurement content is justified according to the age of each participant. As shown in Table 1, the number of psychological behaviour measurements (related to questionnaires and computer-mediated tasks/tests) increases with age. (c,d) Age and sex distributions for participants’ completion in the CKG and PEK Samples (female, red; male, blue). Dots indicate the specific age of each wave’s data collection, while lines indicate the actual intervals between two waves. (e,g) Numbers of participants enrolled (Wave 1) in each age group are calculated according to sex. (f) The actual intervals in the CKG Sample better adhere to the original design; the largest interval is 19 months. (h) In the PEK Sample, intervals have been commonly extended from 16 to 50 months.

### Recruitment Strategy

The devCCNP project focused on enrolling typically developing school-age Chinese children and adolescents. The CKG Sample was included one primary school and one junior high school in Chongqing. The participants were recruited through face-to-face communications between parents, schools, and CCNP program staff. In the case of the PEK Sample, recruitment took place in Beijing, where community-based recruitment was initially accomplished through various science popularization activities and online advertisements. We provided a series of activities for the families to experience educational neuroscience, including lectures on the brain, neuroimaging, cognitive neuroscience, and facility tours to experience MRI mock scanning, to make them interested and familiar with the entire procedure. Because the project gradually gained a good reputation, word of mouth recruitment became a major source of participants.

### Retention Strategy

To accommodate each participant’s after-school schedule, the experimental procedures for one wave were conducted in 2 to 4 separate visits as shown in Table 2. A 1-month time window was given for completing all the experimental protocols in one wave, allowing for flexibility in scheduling. During the COVID-19 pandemic, relevant to the PEK Sample only, the time window was extended to three months to ensure that participants were able to complete the study. In addition, we offered modest monetary compensation and a variety of educational toys to the participants. The primary strategies to promote retention are listed below.

**Table 2.**
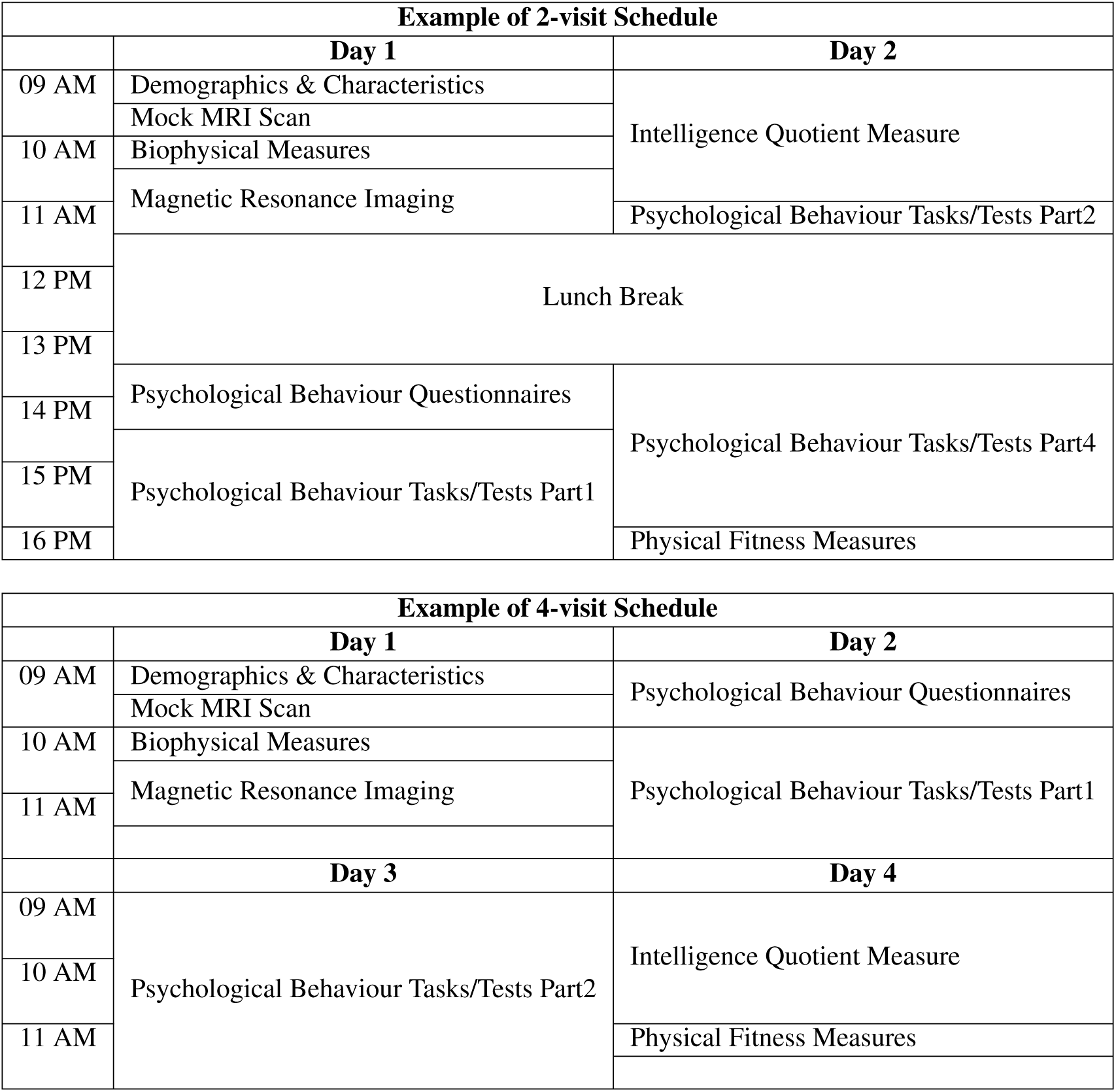
Examples of two individual schedule for each wave’s data collection.

#### Personal Development Report

After each wave’s data collection, every participant was provided with a well-designed personal development report containing feedback on various aspects of physiological characteristics (e.g. height, weight, blood pressure, and heart rate), cognitive ability (e.g. intelligence quotient or IQ), social-emotional development (e.g. social anxiety, depression, stress perception and behavioural problems), personality, and brain development. The brain development report included measurements of global and network morphology (i.e., 7 large-scale brain network organizations^22^). Additionally, the report compared 2 or 3 wave performances to highlight development changes over time. Percentiles or norm-referenced scores were given to guide the interpretation of developmental behaviours. Practical advice or recommendations for enhancing the performance were provided only for reference.

#### Brain Science Popularization

The enrolled participants and their guardians were regularly invited to attend talks popularizing brain science organized by the program staff. The talks were focused on providing an intuitive understanding of the personal development report and promoting extensive knowledge of brain science. During the progress, we emphasized the scientific significance of establishing longitudinal datasets for Chinese children and adolescents, with the aim of encouraging retention in the project. A featured program *Localization of Frontiers for Young Minds articles* (https://kids.frontiersin.org/articles) was launched in July 2019 with weekly neuroscience popularization articles promoted through various social media platforms, such as the *WeChat* Official Accounts Platform. Teenagers volunteered to be part of the translation team and were supervised by the CCNP Science Mentors. This initiative widely popularized background knowledge to school-age students, and improved acceptance of the project among the target population.

### Participant Procedure

#### Screening & Registration

A prescreening phone interview inquired about each participant’s health history, family history of disease, and any potential risk or side effect associated with the MRI procedure. After a detailed introduction, any necessary explanations and an assessment of those who were willing to participate, individuals meeting inclusion criteria without any reason for exclusion were invited to preregistration. Both participants and their guardians were invited to be confirmed on site, and signed the informed consent form before official participation.

The inclusion criteria were as follows:

- Male or female native Chinese speakers aged 6.0–17.9 years at enrolment. Note that some participants under 6 years old were also enrolled as a preexperiment on younger individuals.
- Must have the capacity to provide assent, guardian must have the capacity to sign informed consent.

The exclusion criteria were as follows:

- Guardians unable to provide developmental and/or biological family histories (e.g., some instances of adoption).
- Serious neurological (specific or focal) disorders.
- History of significant traumatic brain injury.
- History or family history (first-degree relatives) of neuropsychiatric disorders, such as ASD, ADHD, bipolar disorder, or schizophrenia.
- Contraindication for MRI scanning, such as metal implants, or pacemakers.

#### Ethical Approval

This project was approved by the Institutional Review Board of the Institute of Psychology, Chinese Academy of Sciences. Prior to conducting the research, written informed consent was obtained from one of the participants’ legal guardians, and written assent was obtained from the participants. Participants who became adults in the longitudinal follow-up provided written consent once becoming 18 years old.

### Experimental Design

Detailed assessments are listed in Table 1. Data collection was accomplished by well-trained research assistants.

#### Demographics & Characteristics

Demographic information (e.g., age, sex and handedness) and characteristics of both participants (e.g., educational level) and their families (e.g., number of children) were collected at the beginning of each wave through self-designed parental questionnaires. The hand preference of the participant was assessed by the Annett Hand Preference Questionnaire (AHPQ)^23^ in the CGK Sample and was classified into 5 subgroups: strong right preference (RR), mixed with right tendencies (MR), mixed (M), mixed with left tendencies (ML), and strong left preference (LL). In the PEK Sample, the Chinese version of Edinburgh Handedness Inventory (EHI)^24^ was applied and participants were classified into 7 subgroups; two additional subgroups compared with the CKG Sample were right preference (R) and left preference (L). Parent-reported Child Behavior Check List (CBCL)^25,26^ was applied with Version: Ages 4–16 (1991 version) in the CKG Sample and Version: Ages 6–18 (2001 version) in PEK Sample. To capture participants’ family characteristics to achieve better population classification, a self-designed parent-reported Subjective Social Status questionnaire using a 10-point self-anchoring scale was additionally conducted in the process of PEK sampling. Additionally, in the PEK Sample, Music Training History Questionnaire for Children^27,28^ was completed by parents to collect information about the participants’ previous training or acquisition of music-related knowledge/skills.

#### Biophysical Measures

Objective biophysical measurements include height, weight, head circumference, and biomarkers of cardiovascular health (i.e., blood pressure and heart rate). The blood pressure assessment was performed immediately after the participant’s MRI scan, and the data provided were related to this specific time point. Visual acuity (naked eyesight in general, corrected eyesight as optional if the participant had ametropia) and Pure Tone Audiometry (PTA)^29^ were specifically measured in the PEK Sample. Even though PTA is a relatively basic and important hearing test, and was conducted in a sound-proof room, we note that the results might be affected by other factors, such as the psychological status of the participant. Therefore, we emphasized that the participant’s biophysical characteristics were only related to the physical and emotional state of the moment.

#### Physical Fitness Measures

Grip strength^30^, standing broad jump^31^ and 15-metre shuttle run^32^ were tested to measure the muscle strength and cardiopulmonary endurance of the participants. After watching the procedure demonstrations, the test method and details were explained to the participants, and they were required to warm up sufficiently. The 15-metre shuttle run was conducted at the end, and the number of completed laps was recorded as the result. Rating of Perceived Exertion (RPE)^33^ was measured immediately after the shuttle run to evaluate exercise intensity.

#### Intelligence Quotient Measure

All participants aged 6–17.9 were given the Wechsler Intelligence Scale for Children-IV-Chinese Version (WISC-IV)^34^ during each wave’s assessment. Ten core subtests and 4 supplementary subtests were combined to estimate Full Scale Intelligence Quotient (FSIQ) with 4 indices: Verbal Comprehension Index (VCI), Perceptual Reasoning Index (PRI), Working Memory Index (WMI) and Processing Speed Index (PSI). Participants aged above 18 years completed the Chinese Version of Wechsler Adult Intelligence Scale (WAIS-IV)^35^.

#### Psychological Behaviour Questionnaires

Widely used questionnaires with high reliability and validity, primarily focused on cognition, personality, and issues pertaining to social-emotional functioning (e.g., life events, self-concept, emotions and affects such as stress, anxiety, depression, loneliness, and positive and negative affect) were obtained by one-on-one instruction. All the psychological behaviour questionnaires corresponding to each Sample are detailed in Table 1.

#### Psychological Behaviour Tasks/Tests

Various experimental paradigms through E-Prime, MATLAB and other platforms were used to assess participants’ cognitive performance in different domains (e.g., executive attention, social cognition, decision-making and language). Some culturally specific tasks were also conducted (e.g., Chinese Character Naming Task). Details are listed in Table 1. Before each formal task/test, participants were informed of the overall procedure through an instructional message and allowed to have exercise trials. Brief introductions on these tasks/tests are as follows:

- **Attention Network Test** The classic Attention Network Test (ANT)^36^ was applied to assess the three attention networks: alerting, orienting and executive attention. During the experiment, small cartoon images of "fish" were presented on the centre of the computer screen for a very short time. Participants were asked to determine as soon and as correctly as possible whether the head of the centre "fish" pointed the left or right (we use images of cartoon fish to replace the "arrow" in the classic ANT paradigm for high preference in children and adolescents). Response times (RTs) and accuracy were measured for each trial.
- **Singleton Stroop Task** This task was introduced to assess an individual’s bottom-up attention capture and top-down inhibitory control. A fixation point was presented on the screen at the beginning and end of each experimental trial. Five short vertical lines were then presented on the screen, and one of the lines was red while the remaining were black. The next task stimulus, a vertical arrow, randomly appeared at the top or bottom of the screen. Participants were asked to respond to the direction of the arrow as quickly and correctly as possible. RTs and accuracy were measured for each trial.
- **Task-Switch Paradigm** In this experiment^37^ participants were asked to make judgements as soon and as correctly as possible between two different types of digit categorization: whether the presented digit was greater or less than 5 and whether the present digit was odd or even. RTs and accuracy were measured for each trial.
- **Digit** *N***-back Task** This paradigm^38^ was used with two levels: 1-back and 2-back. Participants were asked to judge as soon and as correctly as possible whether each stimulus in a sequence, which consisted of nine random digits from 1 to 9, matched the stimulus that appeared *N* items ago. To be specific, participants would determine whether the currently presented digit was the same as the one (i.e., 1-back) or second one (i.e., 2-back) presented before. RTs and accuracy were measured for each trial.
- **Prisoner’s Dilemma** This task was conducted to assess the influence of networks on the emergence of cooperation^39,40^. Before the formal experiment, participants were instructed that there were four blocks of games. Two of them are social partner blocks, in which their partners in each round are peer children and would be paid according to the final outcome. In contrast, in two blocks of nonsocial partner blocks, the partner’s choice was randomly given by computer. In the experiment, first, a fixation point was presented on the screen. Then, a payoff matrix that lists the payoff when the participant and the partner choose to "cooperate" or "betray" is created. Participants were instructed, "You need to choose "cooperate" or "betray" without knowing your partner’s choice. You will then present your partner’s choice and therefore respective benefits based on bilateral choices." Finally, participants were asked to assess their emotional response towards the choices. Before participating in the social decision-making study, both prisoner’s dilemma and ultimatum game introduced next, the participants were asked to describe themselves in a self-introduction, including their age, upbringing, education, personality, and hobbies. The participants were informed that their self-introduction would be anonymously presented to a group of peers who would participate in the same experiments. Those peers acted as their partners in the experiment. Each of those peer partners independently made a choice after reading the participants’ self-introduction and their choices were preprogrammed in the experiment computer and displayed to them in the experiment.
- **Ultimatum Game** This task was designed to explore whether and how social comparisons with third parties affect individual preferences for fair decision-making^41,42^. Before the formal experiment, participants were told that there are two blocks of games. One of them would be under the "gain" context, which means that players in the game are together to distribute gain. The other would be under the "loss" context, which means that players in the game are together to distribute loss suffering. In the formal experiment, an allocation of gain/loss would be offered, and participants needed to decide to accept or reject the offer and report how satisfied they felt about their final rewards/suffering.
- **Delay Discounting Task** To explore reward evaluation and impulsivity characteristics^43^, in this task, participants were asked to make a series of choices to receive a certain value of fictitious funds immediately, or to wait for a period of time (i.e., a day, a week, a month, three months, or six months) before receiving a larger amount. For example, choosing between "Get ¥100 tomorrow" and "Get ¥50 today". The reward amount was presented on the screen immediately after each decision was made.
- **Risky Decision Task** This task was designed as an interactive, sequential gambling game to probe the neural correlates of risk taking and risk avoidance during sensation seeking^44,45^. Participants were instructed to play a roulette game with a certain amount principal at the beginning. After deciding whether to participate in the gamble or not depending on the situation introduced (the odds of winning the jeton), rewards (gain or loss) were presented. Each decision had to be made in 4 seconds. After each trial participants were asked to evaluate and report whether they had made the right choice.
- **Chinese Character Reading Test: Chinese Character Naming Task** This task was introduced to examine children’s reading ability and to determine potential developmental issues in the process of reading acquisition^46^. Participants under 12 years old were asked to read a list of 150 Chinese characters (increasing difficulty from front to back) one by one. The score was calculated from the number of characters reading correctly.
- **Lexical Identification** This task used the semantic priming paradigm to examine mental representations of word meanings and their relationships^47,48^. Critical words consisted of real word targets following a thematic prime (e.g., eat-lunch) or a categorical prime (e.g., apple-banana). Additionally, filler words consisting of nonword targets (e.g., eat-unch) were also added. The words (both the prime and target words) were consecutively presented on the screen and after the presentation of each word, participants were asked to judge whether the word was a real word or not. RTs and accuracy were recorded.
- **Audiovisual Integration of Words** This task examined the integration of visual and auditory word information^49,50^. In each trial, participants were visually presented with one Chinese character on the screen and presented with a word pronunciation at the same time. The character was either audiovisually congruent (where the character and the pronunciation were matching) or incongruent (where the character and the pronunciation were nonmatching). Participants were instructed to judge whether the auditory word pronunciation matched the visual word form. RTs and accuracy were recorded.
- **Brief Affect Recognition Test** This test was used to evaluate an individual’s recognition of facial expressions^51,52^. Participants were first presented with a 200*ms* fixation point in the centre of the screen, and then randomly presented with a picture of a model’s emotional expression for 200*ms*. Ten models (six women and four men) were selected from the Ekman database. Participants were asked to judge the expression presented from two options within the limited time (200*ms*), or they would automatically skip to the next image. Failed to select an expression was marked as wrong. There were 30 sets of facial expressions made up of six different facial emotions (happiness, sadness, fear, disgust, surprise, and anger).
- **Temporal Bisection Paradigm** To evaluate an individual’s characteristics on time perception^53,54^, participants were required to learn two time intervals to strengthen their memory of long and short time intervals. These time intervals were defined as the presenting a 2*cm* × 2*cm* black squares was presented. For short duration, the black squares were presented for 400*ms*, for long duration, the black squares were presented for 1600*ms*. After the training procedure, participants were instructed to judge the length of the test time intervals (rating intervals as "long" or "short") according to the previously learned time intervals. Black squares were randomly presented 20 times for 400, 600, 800, 1000, 1200, 1400, or 1600*ms* interval.
- **Ebbinghaus Illusion** To assess participants’ susceptibility to perceptual illusions^55,56^, participants were instructed to view a screen with a grey background. A probe circle and a reference circle were presented on the left and right sides of the central fixation point. The probe circle was always surrounded by a group of smaller circles. The reference circle, which was fixed in size, was surrounded by larger circles. The perceptual sizes of the probe circle and reference circle were not the same. The task was to adjust the size of the probe circle with up or down arrow key to match that of the reference circle. A chinrest was used to help minimize head movement.
- **Binocular Rivalry** To evaluate sensory eye dominance^57,58^, participants were instructed to view two orthogonal sinewave grating disks (±45^°^ from vertical) dichoptically through a pair of shutter Goggles (NVIDIA 3D Vision2 glasses). A chinrest was used to minimize head motion. The gratings were displayed in the centre of the visual field and were surrounded by a checkerboard frame that promoted stable binocular alignment. Participants were required to report whether they perceived one of the two gratings or non-oriented disks by holding down one of the three keys (Left, Right, or Down arrows) on the keyboard. If a key was not pressed within a predetermined period of time, there would be an audible alarm for the participants.
- **Ocular-tracking Task** This task examines basic visuomotor ability by measuring ocular-tracking performance, as previously described^59–63^. This task was based on the classic Rashbass step-ramp paradigm^64^ modified to accommodate a random sampling of the polar angles from 2^°^ to 358^°^ in 4^°^ increments around the clock face without replacement using 90 trials. Each trial began with a cartoon character (Donald Duck or Daisy, 0.64^°^*H* × 0.64^°^*V*) in the centre of a black background on a computer screen. Participants were asked to fixate on the central character and initiated the trial by pressing a mouse button. After a random delay drawn from a truncated exponential distribution (mean: 700*ms*; minimum: 200*ms*; maximum: 5, 000*ms*), the character would jump in the range of 3.2^°^ to 4.8^°^ away from the fixation point and immediately move back at a constant speed randomly sampled from 16^°^*/s* to 24^°^*/s* towards the centre of the screen and then onwards for a random amount of time from 700 to 1, 000*ms* before disappearing. To minimize the likelihood of an initial catch-up saccade, the character always crossed the centre of the screen at 200*ms* after its motion onset. Both the character speed and moving direction were randomly sampled to minimize expectation effects. Participants were instructed to keep their eyes on the character without blinking once they initiated the trial and then to use their eyes to track the character’s motion as best as they could until it disappeared on the screen.
- **Dichotic Digit Test** This test was used to assess individuals’ binaural integration^65,66^, attention allocation, and auditory/speech working memory ability. A different set of digits (2 or 3 digits) was presented simultaneously to the participant’s left and right ears with an output intensity set to 50*dB* HL. Participants were asked to listen carefully and repeat the digits heard from right ear to left ear during half of the trials, and from left ear to right ear during the other half of the trials. The orders were counterbalanced between participants.
- **Competing Sentences** This test was introduced to examine auditory selective attention and the ability to inhibit irrelevant utterance interference during speech recognition^66^. Two simple Chinese sentences with the same syntactic structure but different contents (7 words with 4 key words, e.g., "the turtle/swims slower/than/the whale" ("乌龟/比/鲸鱼/游得慢")) were presented simultaneously to the left and right ears. Participants were asked to listen carefully and repeat the content in the attended ear (i.e., the output intensity of the attended ear was 35*dB* HL while the nonattended side was 50*dB* HL) at the end of the sentence. The attended ear was left on half of the trial and right on the other half. The orders were counterbalanced between participants.
- **Mandarin Hearing in Noise Test for Children** This test was used to assess speech recognition ability in a noisy environment^67^. A simple Chinese target sentence (15 sentences of 10 Chinese syllables each, e.g., "He drew a tiger with a brush" ("他用画笔画了一只老虎")) was presented by the frontal speaker, and speech spectrum noise was simultaneously played by the frontal or lateral (90^°^ apart) speaker. The noise intensity was constant at 65*dB* SPL, and the starting signal-to-noise ratio (SNR) was 0 dB for the front noise speaker and -5 dB for the side noise speaker. Participants were required to listen carefully and repeat the sentence at the end. The SNR threshold at which participants correctly reported 50% of syllables in the sentence was recorded as the speech recognition threshold (SRT).
- **Verbal Fluency** The verbal fluency test was used to evaluate strategic search and retrieval processes from the lexicon and semantic memory^68,69^. Participants in each trial were required to speak nonrepeated words based on one given category within 1 minute. There were two trials for semantic fluency and two for phonemic fluency. Semantic fluency required participants to say as many words as possible belonging to a particular semantic category (fruit, animal). Phonemic fluency required the participants to say as many different words as possible (excluding proper names) beginning with a Mandarin initial consonant (/d/ and /y/) but not repeating the first vowel and tone. The last four behaviour tests were performed in a soundproof room in one session, and normal hearing in both ears (average hearing threshold *≤* 20*dB* HL from 250 to 8000*Hz*) was required.

#### MRI Mock Scan

In the preparation stage for MRI scans during PEK sampling, mock scanning was performed to improve participant compliance by alleviating anxiety and psychological distress, and to facilitate the success of scans, especially for participants under 12 years old (i.e., primary education stage)^70^. The mock scanner room was built in a child-friendly atmosphere (e.g., child-style decorations, toys or books for different ages, etc) which provided a relaxed buffer zone. A real-size mock scanner built by PST (Psychology Software Tools, Inc.) using a 1:1 model of the GE MR750 3T MRI scanner in use at the PEK site, allowed participants an experience faithful to the actual MRI scanning procedure. Participants were guided to lie still on the bed listening to the recorded MRI scanning sounds and watching the screen through the mirror attached to the model head coil. Three imaging scenarios were performed: resting-state fMRI (rfMRI), morphometric MRI and natural stimulus fMRI (ns-fMRI) which refers to the movie-watching state in this sample. Each scenario lasted at least five and a half minutes. The instructions were consistent with the actual MRI scan, except the movie clip played during the natural stimulus was replaced by additional resources. Head motion data were automatically acquired with the MoTrack Head Motion Tracking System (PST-100722).

#### Magnetic Resonance Imaging

MRI data of CKG Sample were collected using a 3.0-T Siemens Trio MRI scanner (sequencing order: rfMRI*→*T1-weighted*→*rfMRI*→*T2-tse/tirm) at the Center for Brain Imaging, Southwest University. The PEK Sample were imaged on a 3.0-T GE Discovery MR750 scanner at the Magnetic Resonance Imaging Research Center of the Institute of Psychology, Chinese Academy of Sciences (sequencing order: rfMRI*→*T1-weighted*→*rfMRI*→*T2-weighted*→*ns-fMRI*→*DTI). Imaging sequences remained the same across all waves at each site but were different between the two sites and optimized for similar space and time resolutions. Minimal adjustments to sequencing order would occur as necessary. To avoid introducing cognitive content or emotional states into the resting-state condition, the rfMRI scans were always conducted before movie-watching. The detailed acquisition parameters in both samples are presented in Table 3. MRI procedure was performed within one session, small breaks were allowed and instructions were given before starting each sequence. During data collection there were no software or hardware upgrades that would affect the MRI scanning performance.

- **Resting-state fMRI** Two rfMRI scans with identical (within each Sample) parameters were acquired and separated by a T1-weighted sequence. Participants were asked to keep their eyes fixated on a light crosshair (CKG Sample) or a cartoon image (PEK Sample) on the dark screen, to stay still, and not to think of anything in particular. Noise-cancelling headphones (OptoACTIVE^TM^ Active Noise Control Optical MRI Communication System, Version 3.0) were provided in the PEK Sample rfMRI scan to foster a more comfortable imaging experience.
- **Morphometric MRI** Morphometric imaging consisted of T1-weighted, T2-weighted (PEK Sample only) and T2-tse/tir (CKG Sample only) scans. A T2 scan was performed after two rfMRI scans to evaluate brain lesions and improve cross-registration. For both morphometric scans, participants were asked to keep their eyes closed to rest.
- **Natural Stimulus fMRI** This functional MRI condition was implemented in the PEK Sample only and under a movie-watching state. Movie watching mimics real-world experiences related to the context. Movie watching requires the viewer to constantly integrate perceptual and cognitive processing. Movie-watching helps to reduce head motion and increase participant compliance and, therefore, improve the feasibility of brain-behaviour association studies^71^. At the beginning of PEK sampling, participants were watching an audiovisual movie clip that was self-produced and consisted of 6 public interest advertisements^72^ (Table 4). From August 2020, the movie clip was replaced by an animated film named “Despicable Me”^73^ (6*m*: 06*s* clip, DVD version exact times 1: 02: 09 *−* 1: 08: 15, spanning from the bedtime scene to the getting in a car scene).
- **Diffusion Tensor MRI** This sequence was implemented in the PEK Sample only. During the scans participants were free to decide if they wanted to watch another animation clip or rest. Detailed parameters are presented in Table 3.

**Table 3.** MRI Protocol Parameters.

|  | CKG Sample | PEK Sample |  |
| --- | --- | --- | --- |
| Scanner<br>Head Coil | Siemens Trio Tim, 3.0T<br>12-channel | GE Discovery MR750, 3.0T<br>8-channel |  |
|  | r-fMRI | r-fMRI | ns-fMRI |
| Sequence | EPI | Gradient Echo | Gradient Echo |
| Time Acquisition (min:sec) | 7m:45s | 6m | 6m:06s |
| Slices | 38 | 33 | 33 |
| % FOV Phase | 1 | 1 | 1 |
| Resolution (mm) | $3 \times 3 \times 3$ | $3.5 \times 3.5 \times 4.2$ | $3.5 \times 3.5 \times 4.2$ |
| matrix | $72 \times 72$ | $64 \times 64$ | $64 \times 64$ |
| TR (ms) | 2500 | 2000 | 2000 |
| TE (ms) | 30 | 30 | 30 |
| Flip Angle (°) | 80 | 90 | 90 |
| Notes |  |  |  |
|  | T1-weighted | T1-weighted | DTI |
| Sequence | 3D MPRAGE | 3D SPGR | Spin Echo |
| Time Acquisition (min:sec) | 8m:19s | 4m:41s | 10m:54s |
| Slices | 176 | 176 | 75 |
| % FOV Phase | 1 | 1 | 1 |
| Resolution (mm) | $1 \times 1 \times 1$ | $1 \times 1 \times 1$ | $2 \times 2 \times 2$ |
| matrix | $256 \times 256$ | $256 \times 256$ | $112 \times 112$ |
| TR (ms) | 2600 | 6.7 | 8724 |
| TE (ms) | 3.02 | 2.9 | 81.4 |
| Flip Angle (°) | 8 | 12 | 90 |
| Notes | 900ms TI | 450ms TI | 64 directions, 10 b0 images, b=1000s/mm<sup>2</sup> |

**Table 4.** Summary of the advertisements included in the first movie clip used in PEK Sample. (Table 1 in ref^72^)

| Order | Name | Description |
| --- | --- | --- |
| 1 | Animal Humane Society | Cute cats in an animal welfare facility |
| 2 | The World is Where We Live | Side-by-side clips of the human world and the animal world to show the connection between humans and wildlife |
| 3 | A Love Story in Milk | A love story between two milk bottles |
| 4 | Water ink | Painting with water and black ink |
| 5 | 50 Years of Flower Power | A story about the Struggle for Human Rights in the last 50 years |
| 6 | You can help save the Cerrado | Fingertut demonstrating the wildlife |

### Summarizing lessons learned

Throughout the implementation of the pilot devCCNP, we faced several challenges and gained valuable insights. We are continuing to improve strategies in dissemination, recruitment, retention, and characterization. Here are some key considerations that may aid similar endeavours, including large-scale sampling projects (e.g. the national longitudinal cohort on child brain development in China).

#### Recruitment Strategy

Due to the particularity of children and adolescents, studies involving this population typically encounter significant challenges. All projects should be conducted on the premise of not affecting academic progress and ensuring safety. Both school- and community-based recruitment have distinct advantages and, inevitably, inherent drawbacks.

- **School-based Strategy** Support of flexible schoolwork arrangements matched with the sampling schedule can greatly ensure the quantity and quality of data collection for junior and senior high school participants. With the help and encouragement from coordinators in school, recruitment efforts could be reduced. However, for these same reasons, the participants’ motivation could be compromised, as they may not be primarily driven by their interest in the project or may lack a clear understanding of the value and the contribution of participation. Meanwhile selecting recruiting schools may also, to some extent, reduce the sample representativeness of the target population.
- **Community-based Strategy** Younger participants are undoubtedly easier to recruit in the community, but the number of pubertal-age participants is limited especially for longitudinal studies. Self-enrolled participants recruited at the community level or their guardian typically possess relevant knowledge and understand the value of participating in the project; therefore they are strongly motivated and tend to cooperate better. However, this also biases the sample to families with higher levels of education, or with some uncertain developmental problems. Especially with word of mouth spreading and popularizing, the similarity between participants’ families (e.g., social status, economic background) and/or their characteristics would be higher, which might diminish the individual differences between the participants.

The drawbacks outlined above can be compensated by combining diverse recruitment strategies and expanding the age range and geographical regions of the recruitment. This approach could enable a greater diversity of physical, psychological and cognitive phenotypes and promote the establishment of a typical developing cohort.

#### Experimental Design

Charting the typical developmental trajectories of individuals (with respect to physical, psychological and morphological development) through longitudinal design greatly contributes to uncovering the complex relationship between the brain and behaviour. As a long-term project, it is important not only to assess the full range of participants’ current state at a single time point of data collection, but also to capture what important life or social events occur during the follow-up period. These events include, but are not limited to, a family event (e.g., death of a family member, divorce of guardians), the birth of siblings, sudden illness, a significant social or public health event, and others. Future projects could employ regular questionnaires or scales (e.g., monthly) to collect related information during follow-up intervals, so that relevant details could be recorded. Alternatively, participants could be asked to retrospectively report the events at each time point of data collection, but this may miss the ability to capture their physiological or psychological experiences at the time of the event.

#### Practical Experience

The attentiveness and compliance of participants have significant impacts on data quality. The following lists the lessons we have learned in the course of our practice.

- **Questionnaire and Scale** For large-scale projects involving different economic or cultural areas (e.g., northern and southern China), it is recommended to apply both questionnaires or scales consisting of subjective and objective assessments. For example, it is suggested to apply Subjective Social Status and to inquire about family income to assess participants’ family economic status. The combination of objective and subjective questions for the same evaluation purpose can better classify populations living in areas of significant cultural differences. This recommendation also applies to other physical, psychological and cognitive assessments.
- **Behaviour Tasks/Tests** Most of the behavioural measurements tested with computers require convenient interactions with participants. Tasks requiring participants to press keys quickly is not conducive to young children if the keys are too small or placed too close. For example, pressing "1" or "2" on a keyboard is more likely to cause errors than pressing "A" or "M". Some measurements have higher requirements with respect to participant posture (e.g., ocular-tracking task requires the participant to operate the mouse while keeping the head and upper body still). Therefore, the number of trials and the duration of each trial need to be carefully designed. An overall time of less than 20 minutes for completion is recommended for young participants. At the same time the related hardware equipment should be able to accommodate a broad range of participant characteristics (e.g., head circumference, height, bodily form). For example, common ocular-tracking devices in the laboratory need to be equipped with stable chairs that can be adjusted for a wide range of heights, child-sized desks, or chinrests that can restrain the head. It is recommended to invite children of each age group to evaluate all the experimental protocols at the design stage.
- **Magnetic Resonance Imaging** A scanning time for one MRI session of no longer than 45 minutes (one hour maximum) is strongly recommended, especially for junior participants. In prticular, mock scan training before formal MRI was shown to effectively improve the success of imaging. In general, training immediately before the formal MRI can be effective although additional mock training episodes before the formal MRI day could also be considered if the participants are particularly scared, are sensitive to sound, or find it difficult to concentrate.
- **Personalized Schedule** During each wave’s data collection, as shown in Table 2, the order in which the tests are scheduled needs to be thoroughly arranged. To better achieve MRI data collection in this project, in principle, MRI was arranged at the beginning of each wave. For those who had more than 2 visits within one wave, IQ measurements were scheduled on the last visit, as they were usually of greater interest to guardians. Physical fitness tests should not be scheduled within a few hours before MRI scans; measurements concerning visual perception should not be scheduled after the measurement that require staring at digital screen for an extended duration.
- **Implementation Progress** Generally, one-on-one instruction from the same implementer across visits (within or even across waves) can be conducive to friendly and cooperative relationships with the participants and can be, especially helpful in relieving the timidity of young children to strangers. Measurements with higher qualification requirements for the implementer (i.e., IQ measure) are recommended to be conducted by limited authorized staff. It is worth mentioning that, unless it is ethically required, we do not recommended that parents be allowed to observe the participant’s engagement process, as this may have the potential to impact their child’s performance.

## Data records

### Dataset Deposition

The devCCNP data has been publicly shared in the *Chinese Color Nest Project (CCNP) – Lifespan Brain-Mind Development Data Community* (https://www.scidb.cn/en/c/ccnp), which is a public platform supported by the National Science Data Bank for sharing CCNP-related data and promoting the cooperation of open neuroscience.

#### devCCNP full release

This dataset will be fully available to the research community when acquisition is completed for the pilot stage of CCNP. Before that stage, the full (both MRI and behaviour) data are only available to researchers and collaborators of CCNP. Of note, all of the MRI data have been deposited into the Science Data Band and are accessible upon requests submitted according to the instructions on the Science Data Bank website (https://doi.org/10.57760/sciencedb.07478). A sample of the longitudinal data from a participant is fully accessible at FigShare (https://doi.org/10.6084/m9.figshare.22323691.v1) to demonstrate the data structure.

#### devCCNP Lite

This release version contains only T1-weighted MRI, rfMRI and diffusion tensor MRI data of devCCNP. No cognitive or behavioural information is included. The devCCNP Lite is immediately accessible upon the requests according to the instructions on the Science Data Bank website (https://doi.org/10.57760/sciencedb.07860).

### Data Structures

All data files are organized according to the Brain Imaging Directory Structure (BIDS) standards^21^. An example of the MRI data storage structure is presented in Figure 2. Under the top-level project folder "devCCNP/", CKG and PEK Sample are organized separately. Each participant’s folder "sub-CCNP*/" may contain several subfolders depending on how many waves have been completed to date (i.e., if all waves are finished, the folder would include three "/ses-*" subfolders). Imaging data (".nii.gz") and metadata (".json") are organized into modality-specific directories "/anat/", "/func/" and "/dwi/". Note that in the PEK Sample, Diffusion Tension Imaging (DTI) data files are stored under "/dwi/" folders with the datatype name "*_dwi.*". All demographic and behavioural data are structured under the "/beh/" folder (".tsv"). Detailed parameters of each psychological behaviour task/test are provided in the "json" file attached.

**Figure 2.**
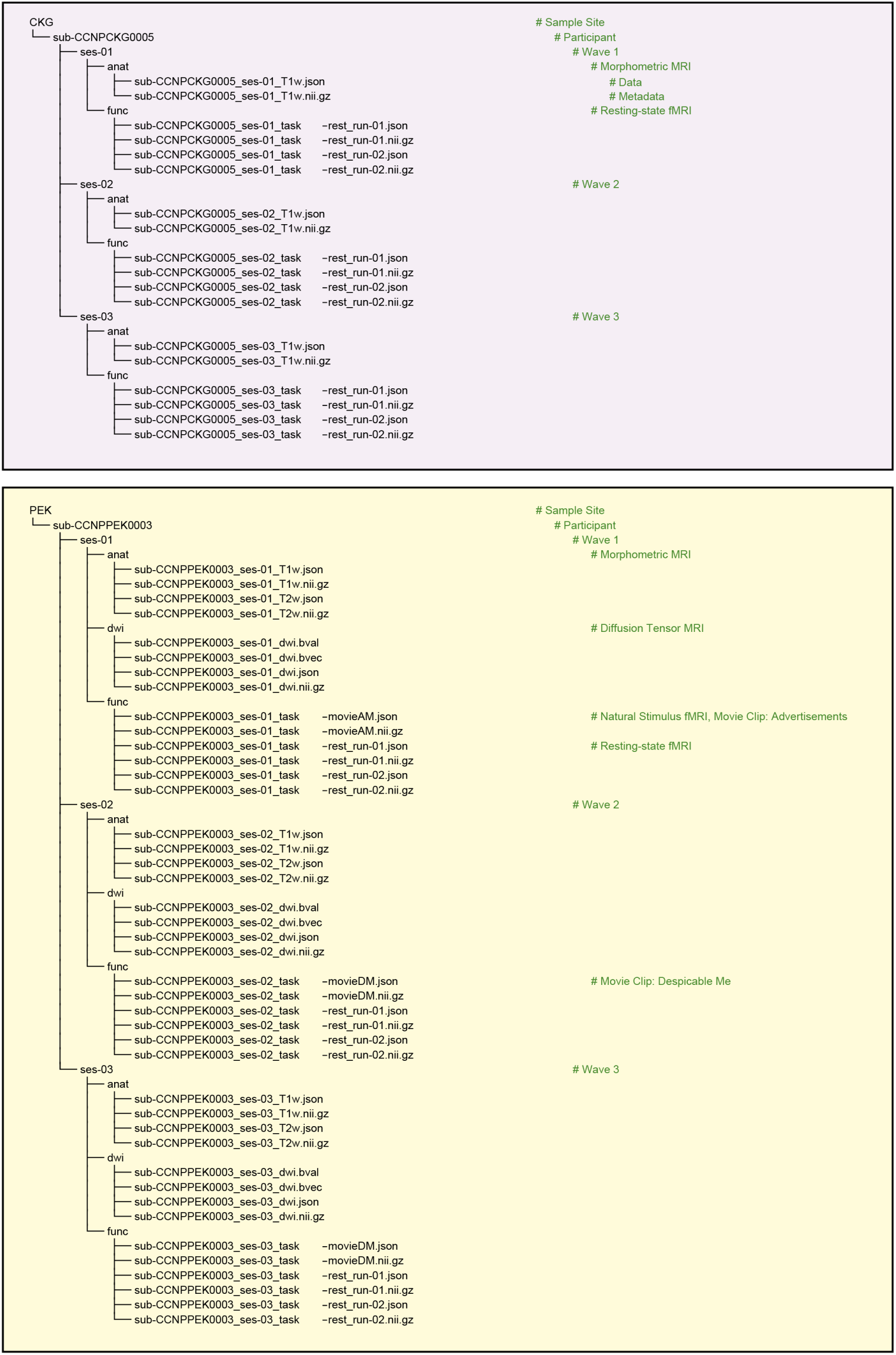
Example of the MRI raw data directory structure. Collected MRI raw data are structured within a hierarchy of folders according to the standard BIDS format. Under the toplevel project folder "devCCNP/", CKG (top) and PEK (bottom) Samples are organized separately. Each participant’s folder "sub-CCNP*/" may contain several subfolders depending on how many waves have been completed to date (i.e., if all waves are completed, the folder would include three "/ses-*" subfolders). Imaging data (".nii.gz") and metadata (".json") are organized into modality-specific directories "/anat/", "/func/" and "/dwi/". Note that in the PEK Sample, Diffusion Tension Imaging (DTI) data files are stored under "/dwi/" folders with datatype name "*_dwi.*".

### Partial and Missing Data

Some participants were not able to complete all components of the CCNP protocol due to a variety of situations (e.g., delay or cancel caused by the COVID-19 pandemic). Overall, we logged data collection if any issues occurred that required extra attention during analysis (see details written in the "json" file attached to each data).

### Data Licence

The devCCNP sample data licence is CC-BY-NC 4.0. To access data, investigators must complete the application file *Data Use Agreement on Chinese Color Nest Project* (DUA-CCNP) located at: http://deepneuro.bnu.edu.cn/?p=163 and have it reviewed and approved by the Chinese Color Nest Consortium (CCNC). Compliance with all terms specified by the DUA-CCNP is required. Meanwhile, the baseline CKG Sample on brain imaging is available to researchers via the International Data-sharing Neuroimaging Initiative (INDI) through the Consortium for Reliability and Reproducibility (CoRR)^74^. More information about CCNP can be found at: http://deepneuro.bnu.edu.cn/?p=163 or https://github.com/zuoxinian/CCNP. Requests for further information and collaboration are encouraged and considered by the CCNC; please read the Data Use Agreement and contact us via.

## Technical Validation

### Sample composition

A total of 479 participants completed baseline visits, 247 (51.6%) completed the second wave data collection, and 138 (28.9%) completed the third wave (i.e., final protocol) as of December 2022. There were 648 (75.0%) measurements completed by participants twelve years old or younger. The number of participants who completed visits in each age cohort are shown in Table 5, and age and sex composition are presented in Figure 1c-d. Demographic and enrolment data for both the CKG Sample (enrolled in 2013-2017) and the PEK (enrolled in 2018-2022) Sample are listed in Table 6. As mentioned above, the overall design has a longitudinal follow-up interval of 15 months, to which the CKG Sample consistently adhered; however, during the PEK sampling, the intervals were prolonged. For instance, inevitable practical situations affected community-based recruitment, primarily the COVID-19 pandemic. Please note that during COVID-19, data collection was suspended from January to August 2020. We designed questionnaires to assess participants’ learning and daily life status^12^. Each participant’s sampling age and corresponding intervals are presented in Figure 1e-h. For all of the measurement intervals, 122 (32.1%) were achieved by design.

**Table 5.** Enrollments of each age cohort in two Samples.

| Age Group | Sample | ≤5 | 6 | 7 | 8 | 9 | 10 | 11 | 12 | 13 | 14 | 15 | 16 | 17 | ≥18 | Enrollments |
| --- | --- | --- | --- | --- | --- | --- | --- | --- | --- | --- | --- | --- | --- | --- | --- | --- |
| Wave1 | CKG | 0 | 7 | 21 | 19 | 22 | 27 | 29 | 10 | 20 | 10 | 10 | 20 | 12 | 0 | 207 |
|  | PEK | 8 | 37 | 51 | 31 | 47 | 36 | 24 | 16 | 5 | 4 | 4 | 3 | 3 | 3 | 272 |
| Wave2 | CKG | 0 | 0 | 2 | 17 | 18 | 24 | 20 | 30 | 7 | 14 | 9 | 7 | 9 | 6 | 163 |
|  | PEK | 0 | 0 | 7 | 12 | 15 | 13 | 17 | 10 | 4 | 4 | 1 | 0 | 0 | 1 | 84 |
| Wave3 | CKG | 0 | 0 | 0 | 0 | 8 | 25 | 15 | 6 | 18 | 7 | 7 | 7 | 2 | 6 | 101 |
|  | PEK | 0 | 0 | 0 | 1 | 2 | 10 | 3 | 8 | 4 | 3 | 4 | 2 | 0 | 0 | 37 |
| Total |  | 8 | 44 | 81 | 80 | 112 | 135 | 108 | 80 | 58 | 42 | 35 | 39 | 26 | 15 | 864 |

**Table 6.**
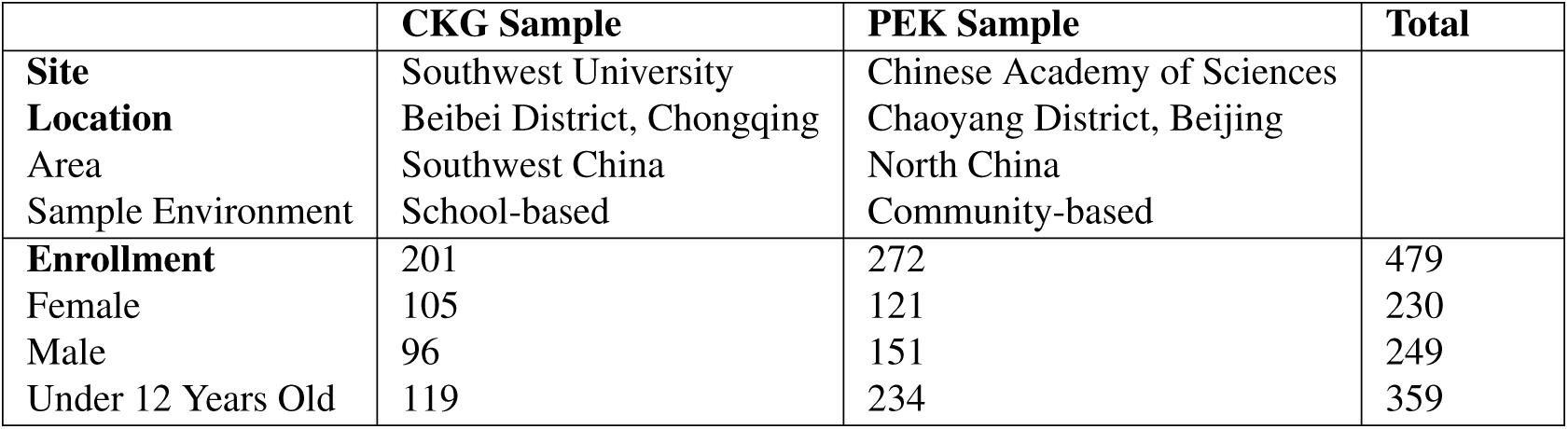
Enrollment profile at two Samples.

|  | **CKG Sample** | **PEK Sample** | **Total** |
| --- | --- | --- | --- |
| **Site** | Southwest University | Chinese Academy of Sciences |  |
| **Location** | Beibei District, Chongqing | Chaoyang District, Beijing |  |
| **Area** | Southwest China | North China |  |
| **Sample Environment** | School-based | Community-based |  |
| **Enrollment** | 201 | 272 | 479 |
| Female | 105 | 121 | 230 |
| Male | 96 | 151 | 249 |
| Under 12 Years Old | 119 | 234 | 359 |

### Quality Assessment

#### Phenotypic Data

All of the psychological and behavioural data were made available to users regardless of data quality. We provided all the information on situations that may affect the quality of data within the "json" file. This can guide investigators decisions regarding inclusion of the result data. To verify whether the measured distributions obey a normal distribution, we performed preliminary statistical analysis of several core behaviour measures in the dataset (Figure 3). Distributions of FSIQ and four indices are shown in Figure 3a. We summarize the median, mean and standard deviation for each Sample. As shown in Table 7, the Shapiro-Wilk test suggests that the sample data commonly disobey a Gaussian distribution. We believe that this is a common situation that arises when recruiting from the local community (PEK sample), as the program tends to attract parents with high levels of education who place greater emphasis on education. Better education conditions could result in higher IQ. Furthermore, the IQ scores were normalized based on the normative model of Chinese children established in 2008^34^, which may be out of time. Additionally, there was a significant difference between the FSIQ, WMI, PRI and VCI performance of the two samples as identified by the rank-sum test. Mental health assessed by CBCL scores demonstrated that the majority of participants were in the normal range (Figure 3b) with only 12 participants (1.77%) exhibiting CBCL total problem scores *≥* 70. We performed preliminary statistical analysis of several core characterization measures and present their accuracy rates in Figure 3c.

**Figure 3.**
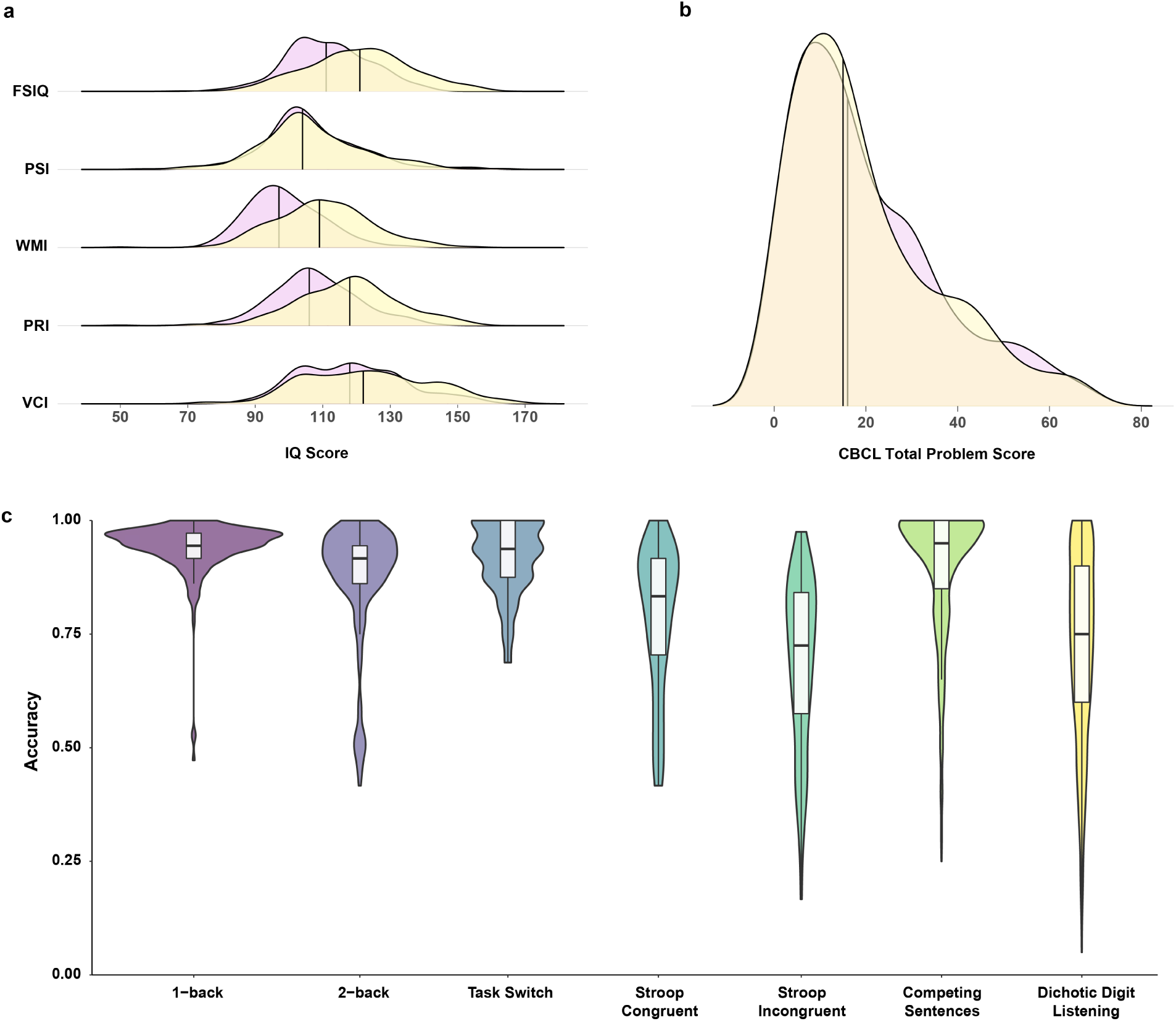
Example of performance on several core characterization measures. (a) Distribution of Full Scale Intelligence Quotient (FSIQ) with four indices: Processing Speed Index (PSI), Working Memory Index (WMI), Perceptual Reasoning Index (PRI), and Verbal Comprehension Index (VCI). Related statistical results are shown in Table 7. (b) Distribution of CBCL total problem scores. Two samples are displayed separately (CKG, light pink; PEK, canary) in (a-b), and vertical lines indicate the medians of samples. (c) Distribution of accuracy rates for seven behaviour measurements. Extremely low values are removed for plotting. Data are represented for measurements of all waves.

**Table 7.**
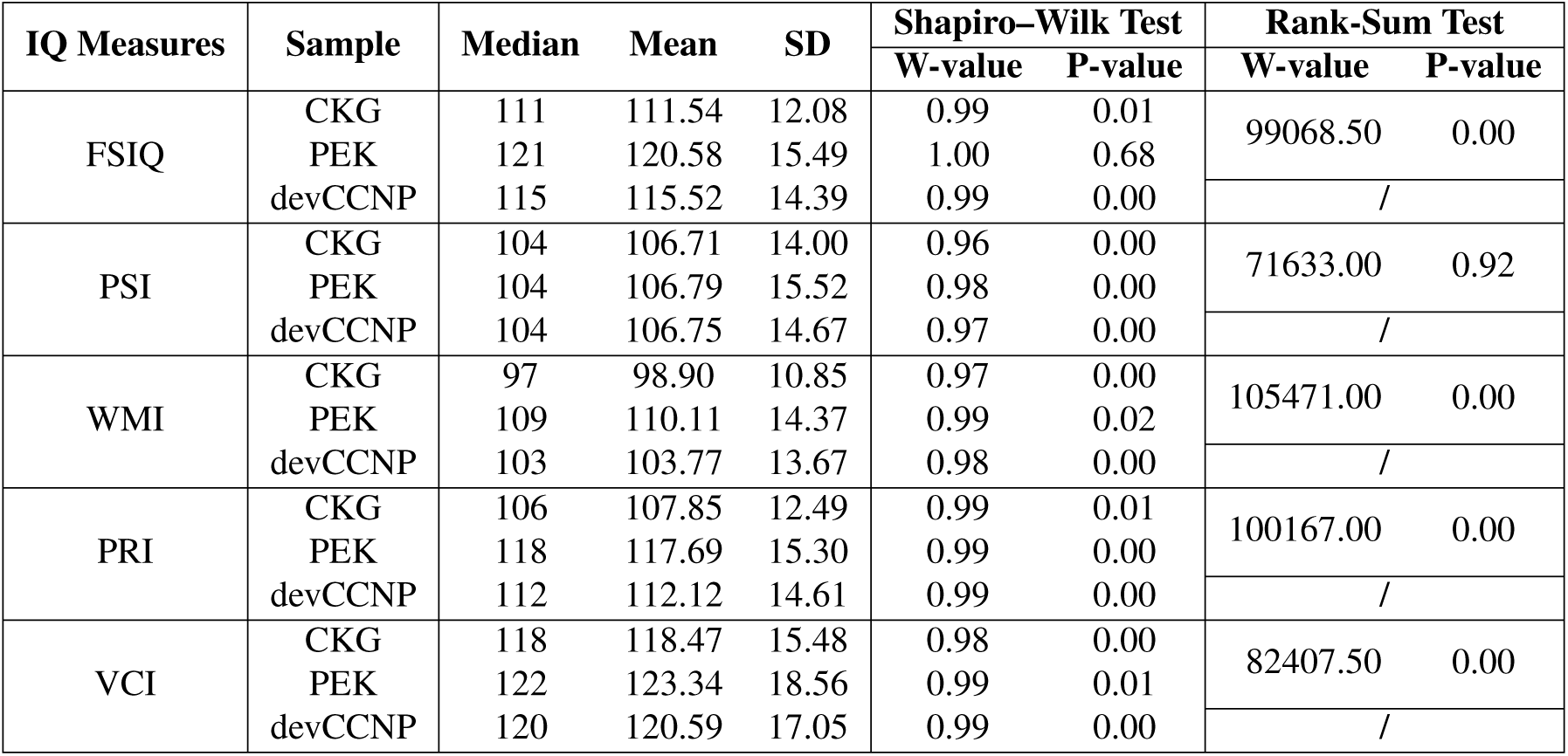
Distribution and statistics of IQ measures.

| IQ Measures | Sample | Median | Mean | SD | Shapiro–Wilk Test |  | Rank-Sum Test |  |
| --- | --- | --- | --- | --- | --- | --- | --- | --- |
|  |  |  |  |  | W-value | P-value | W-value | P-value |
| FSIQ | CKG | 111 | 111.54 | 12.08 | 0.99 | 0.01 | 99068.50 | 0.00 |
|  | PEK | 121 | 120.58 | 15.49 | 1.00 | 0.68 |  |  |
|  | devCCNP | 115 | 115.52 | 14.39 | 0.99 | 0.00 |  |  |
| PSI | CKG | 104 | 106.71 | 14.00 | 0.96 | 0.00 | 71633.00 | 0.92 |
|  | PEK | 104 | 106.79 | 15.52 | 0.98 | 0.00 |  |  |
|  | devCCNP | 104 | 106.75 | 14.67 | 0.97 | 0.00 |  |  |
| WMI | CKG | 97 | 98.90 | 10.85 | 0.97 | 0.00 | 105471.00 | 0.00 |
|  | PEK | 109 | 110.11 | 14.37 | 0.99 | 0.02 |  |  |
|  | devCCNP | 103 | 103.77 | 13.67 | 0.98 | 0.00 |  |  |
| PRI | CKG | 106 | 107.85 | 12.49 | 0.99 | 0.01 | 100167.00 | 0.00 |
|  | PEK | 118 | 117.69 | 15.30 | 0.99 | 0.00 |  |  |
|  | devCCNP | 112 | 112.12 | 14.61 | 0.99 | 0.00 |  |  |
| VCI | CKG | 118 | 118.47 | 15.48 | 0.98 | 0.00 | 82407.50 | 0.00 |
|  | PEK | 122 | 123.34 | 18.56 | 0.99 | 0.01 |  |  |
|  | devCCNP | 120 | 120.59 | 17.05 | 0.99 | 0.00 |  |  |

#### Structural MR Imaging

Structural MRI images were first anonymized to remove all facial information from the raw MRI data. We obscured the facial information using the face-masking tool customized with the Chinese paediatric templates^16^. The anonymized images were then denoised by spatially adaptive nonlocal means and corrected for intensity normalization in the Connectome Computation System (CCS)^75^. To extract individual brains, we trained a deep learning method using a small set of semiautomatically extracted brains in the CKG Sample, and then applied it to all the devCCNP samples. The preprocessed brain volumes were all in the native space and fed into the FreeSurfer (version 6.0) pipeline to obtain general morphological measurements of different brain morphometry. We visually inspected the quality of the T1-weighted images, and two raters were trained to rate the quality using a 3-class framework^76^, with "0" denoting images that suffered from gross artefacts and were considered unusable, "1" with some artefacts, but that were still considered usable, and "2" free from visible artefacts. Images with an average score lower than "1" across the two raters were excluded. A total of 761 (91.9%) images passed the quality control, with 436 (94.8%) images in the CKG Sample and 325 (88.3%) images in the PEK Sample. The Spearman’s rank correlation coefficient of the two raters was 0.495.

#### Functional MR Imaging

RfMRI data preprocessing^75^ included the following steps: (1) dropping the first 10*s* (5 TRs) for the equilibrium of the magnetic field; (2) correcting head motion; (3) slice timing; (4) despiking for the time series; (5) estimating head motion parameters; (6) aligning functional images to high resolution T1 images using boundary-based registration; (7) mitigating nuisance effects such as ICA-AROMA-derived, CSF and white matter signals; (8) removing linear and quadratic trends of the time series; and (9) projecting volumetric time series to *fsaverage5* cortical surface space. Scans with a mean FD greater than 0.5 were excluded. A total of 452 (98.3%) scans in the CKG Sample and 328 (92.4%) scans in the PEK Sample had at least one rfMRI passed the quality control in each session.

### Brain Growth Charts

Growth charts on height, weight and head circumference are a cornerstone of paediatric health care. A similar tool has been recently generated for lifespan development of human brain morphology^18^ by the Lifespan Brain Chart Consortium (LBCC) (https://github.com/brainchart/lifespan). While promising for characterizing the neurodevelopmental milestones and neuropsychiatric disorders, these charts need more diverse samples to enhance their utility in practice. Here, we employed the devCCNP Sample and the NKI-Rockland Sample (NKI-RS) for Longitudinal Discovery of Brain Development Trajectories^77^ to upgrade the LBCC charts. All the preprocessed T1-weighted MRI images from devCCNP and NKI-RS were subjected to the same manual quality control procedure from the same raters at each site.

Specifically, the maximum likelihood method was used to estimate sample-specific or site-specific statistical offsets (random effects, i.e., mean *µ*, variance *σ*, and skewness *υ*) from the age- and sex-appropriate epoch of the normative brain growth trajectory modelling through the Generalized Additive Models for Location, Scale and Shape (GAMLSS: see details of the site-specific growth chart modeling in Figure 5 from the LBCC original work^18^). Out-of-sample centile scores for each participant from the devCCNP and the NKI-RS site benchmarked against the offset trajectory were estimated. The normative growth trajectories were estimated for not only global neurotypes including total cortical grey matter volume (GMV), total white matter volume (WMV), total subcortical grey matter volume (sGMV), global mean cortical thickness (CT) and total surface area (SA) but also, regional neurotypes, including volumes of the 34 neuroanatomical areas according to the Desikon-Killiany (DK) parcellation^78^.

According to the lifespan WMV trajectory from the LBCC seminal work^18^ (i.e., rapid growth from mid-gestation to early childhood, peaking in young adulthood at 28.7 years), we presented the growth curves of WMV for devCCNP-CKG, deveCCNP-PEK and NKI-RS (Figure 4). These curves indicated rapid increases in WMV from childhood to adolescence consistent with the LBCC findings. To better illustrate the growth curve differences between populations, we depicted site- and sex-specific (adjusted) growth curves of WMV in Figure 4 (top). WMV is made up of the connections between neurons for cortical communications via neural information flow, and thus, its growth reflects underlying microstructural plasticity during school-age neurodevelopment^79^ (e.g., language performance and training effects during learning^80^). In our analyses, the study-specific variability (e.g., imaging or sample bias) was adjusted by the GAMLSS modelling method. Therefore, the findings we detected are more reproducible and generalizable across devCCNP and NKI-RS samples. Specifically, as shown in Figure 4, boys had larger WMV than girls, whereas the CKG participants (bottom, right) exhibited relatively smaller WMV than the participants from PEK (bottom, middle) and NKI-RS (bottom, left). Brain growth curves are included in the Supplementary Information (GMV, Figure S1; sGMV, Figure S2; TCV, Figure S3; mean CT, Figure S4; TSA, Figure S5).

**Figure 4.**
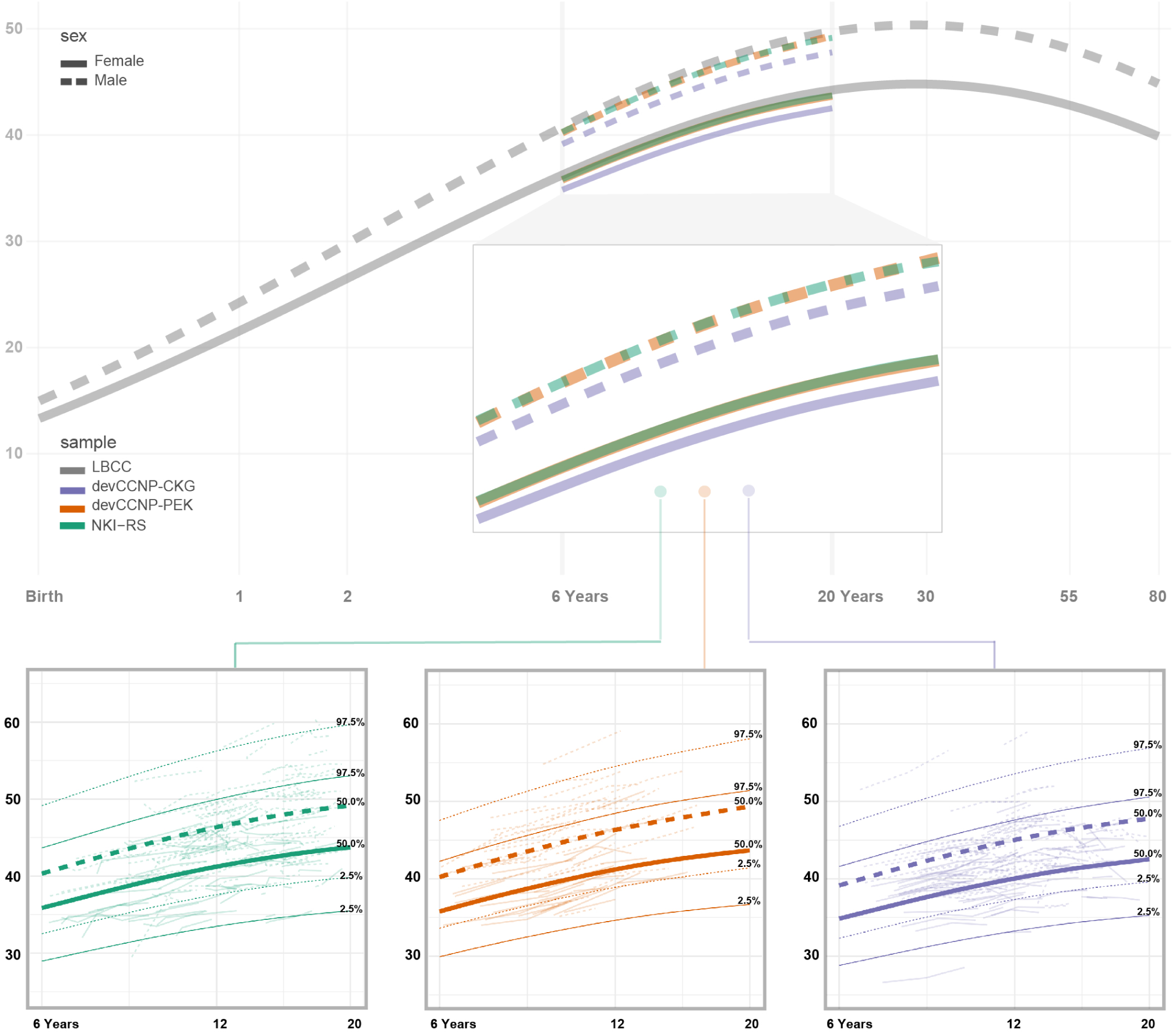
Site/sex-specific brain charts of white matter volume (WMV). The sex-specific lifespan brain charts of WMV (LBCC, light gray) were adjusted by leveraging the school-aged (6–18 years old) samples for three sites (devCCNP-CKG, purple; devCCNP-PEK, orange; NKI-RS, green). The site-specific brain charts are depicted with their percentiles (2.5%, 50%, 97.5%) for males (dashed lines) and females (solid lines). The background polylines characterize individual WMV changes (unit: 10*ml* or 10, 000*mm*^3^) extracted from the multicohort accelerated longitudinal samples.

To quantitatively estimate the diversity in brain growth attributable to ethnicity (referring between devCCNP to NKI-RS) and geographics (referring between devCCNP-CKG to devCCNP-PEK), we computed the normalized variance (NV)^16^ of regional volume for each DK-parcel with the following equation

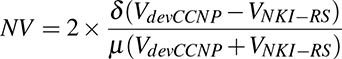

, where *V* is a vector referring to the parcel volume and *δ* referrs to the standard deviation. In other words, NV indicates the degree of curve shape dispersion between two growth curves across ages (we use 0.1 year as the sample age). *δ* is normalized by the mean volume of parcels across two samples, denoted as *µ*. The results are illustrated in Figure 5 (first row). A small NV indicates that two brain growth curves share similar shapes, and vice versa. The sex-specific lifespan brain charts of regional volume (unit: *ml*) specific to one high NV (Pars Orbitalis; second row) and one low NV (Paracentral Lobule; third row) were depicted to illustrate the differences. As shown in Figure 5 (bottom), we matched the 34 parcellated regions to the 8 large-scale functional networks^81^ for an intuitive sense of the growth chart differences at the network level. We present the NV rank for comparisons between NKI-RS and devCCNP as well as between devCCNP-CKG and devCCNP-PEK in Figure 5 (forth row). As done in the LBCC paper^18^, we built normative growth charts of a brain parcel by GAMLSS modelling on the total volume of the parcel as the sum of its two homotopic areas in the two hemispheres. The NV and its rank maps were rendered onto both lateral and medial cortical surfaces of the left hemisphere for visualization purposes. Details of NV are listed in Table S1 and S2 according to their ranking orders. Individual differences in growth charts of cortical volumes between devCCNP and NKIR-RS are much larger than those between CKG and PEK. Such differences are spatially ranked in a consistent order among populations, indicating more diverse growth curves among individuals in high-order associative (frontoparietal or cognitive control, ventral attention, default mode and language) areas than those in primary areas.

**Figure 5.**
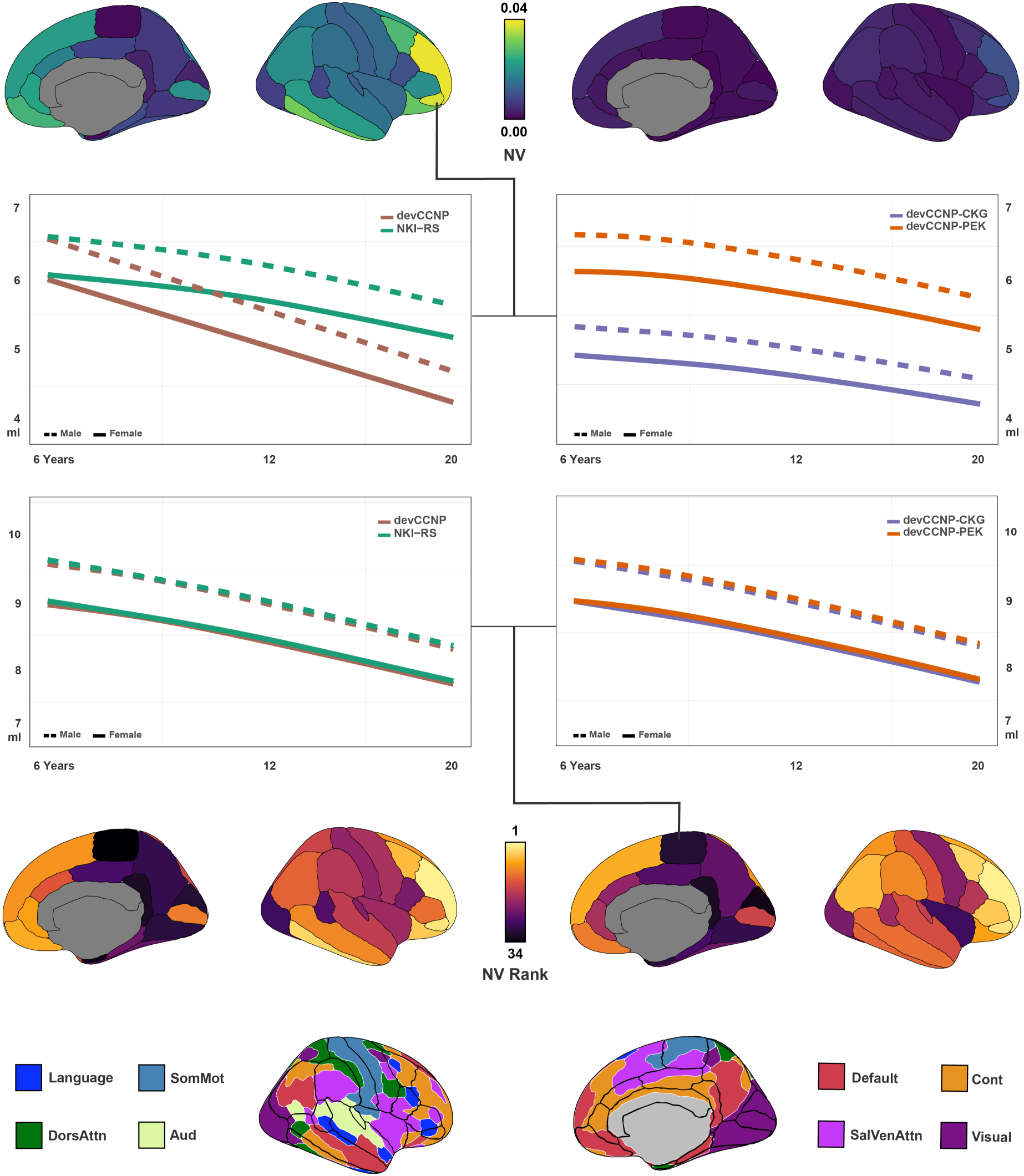
Similarities in brain growth curves between devCCNP and NKI-RS. NV values of the similarity between the United States and China (first row, left) and two samples within devCCNP (first row, right) are presented through 34 gyral-based neuroanatomical regions, referred to as Desikon-Killiany parcellation^78^ (bottom, matched to Kong2022 8 large-scale functional network order^81^). Sex-specific lifespan brain charts of regional volume (unit: *ml*) specific to one high NV (pars orbitalis; second row) and one low NV (paracentral lobule; third row) are depicted. Males are denoted with dashed lines and females are denoted with solid lines. On this basis, the ranks of NV values of these regions are presented (forth row) from highest to lowest. See Table S1 and S2 for detailed values. Note that only the NV values and rank of female participants are shown here, as brain charts are modelled sex-specifically^18^. The left hemisphere is plotted here purely for visualization purposes. See Figure S6 for results relating to male participants.

## Usage Notes

Part of this dataset has been successfully used in our previous publications. Two review articles (in Chinese^15^ and English^12^) were published to summarize the devCCNP protocol for experimental design, sample selection, data collection, and preliminary key findings in stages. With the baseline brain imaging data from devCCNP-CKG, we previously reported that children exhibited similar region-specific asymmetry of the dorsal anterior cingulate cortex (dACC) as adults, and further revealed that dACC functional connectivity with the default, frontoparietal and visual networks showed region-specific asymmetry^82^. Head motion data during mock scanning from devCCNP-PEK were used to demonstrate frequency-specific evidence to support motion potentially as a developmental trait in children and adolescents by the development of a neuroinformatic tool DREAM^83^. Social anxiety was positively correlated with the GMV in an area of the orbital-frontal cortex, and its functional connectivity with the amygdala^84^. A standardized protocol for charting brain development in school aged children has been developed to generate the corresponding brain templates and model growth charts, revealing differences in brain morphological growth between Chinese and American populations particularly around puberty^16^. Meanwhile, by manual tracing, we charted the growth curves of the human amygdala across school ages through longitudinal brain imaging^85^. Using rfMRI data, we revealed age-dependent changes in the macroscale organization of the cortex, and the scheduled maturation of functional connectivity gradient shifts, which are critically important for understanding how cognitive and behavioural capabilities are refined across development, marking puberty-related changes^17^.

The baseline imaging data of the CKG Sample has been released as part of the CoRR^74^ and the IPCAS 7 site (http://dx.doi.org/10.15387/fcp_indi.corr.ipcas7), which has been listed as one of the existing, ongoing large-scale devel-opmental dataset^86^. As part of an international consortium recently initiated for the generation of human lifespan brain charts^18^, CCNP contributes to the largest worldwide MRI samples (*N* > 120, 000) for building normative brain charts for the human lifespan (0 *−* 100 years). The full set of devCCNP data is increasingly appreciated by collaborative studies on school-aged children and adolescents. All data obtained freely from the INDI-CoRR-IPCAS7 or CCNC, can only be used for scientific research purposes. The users of this dataset should acknowledge the contributions of the original authors, properly cite the dataset based on the instructions on the Science Data Bank website (https://doi.org/10.57760/sciencedb.07478 and https://doi.org/10.57760/sciencedb.07860). We encourage investigators to use this dataset in publication under the requirement of citing this article and contact us for additional data sharing and cooperation.

## Code availability

No custom codes or algorithms were used to generate or process the data presented in this manuscript.

## Acknowledgements

CCNP receives funding support from the Start-up Funds for Leading Talents at Beijing Normal University, the National Basic Science Data Center “Chinese Data-sharing Warehouse for *In-vivo* Imaging Brain” (NBSDC-DB-15), the Key-Area Research and Development Program of Guangdong Province (2019B030335001), the Beijing Municipal Science and Technology Commission (Z161100002616023, Z181100001518003), the Major Project of National Social Science Foundation of China (20&ZD296), the CAS-NWO Programme (153111KYSB20160020), the Guangxi BaGui Scholarship (201621), the National Basic Research (973) Program (2015CB351702), the Major Fund for International Collaboration of National Natural Science Foundation of China (81220108014), the Chinese Academy of Sciences Key Research Program (KSZD-EW-TZ-002) and the National Basic Research Program (2015CB351702). CCNC wish to thank all the community partners, research participants, and families who took part in this project. We thank additional team members who supported data acquisition and management. We are grateful to all the research assistants for participating in data collection. We thank numerous expert consultants who contributed to the protocol development. We are grateful to data assistance by Dr. Ting Xu from Child Mind Institute and computing resources of storage and processing provided by the Chinese Academy of Sciences and Beijing Normal University.

## Author Information

### Contributions

Conception and Design: X-N.Z., M.B., A.C., X.C., Y.D., T.F., L.L., S.L., X.L., J.Q., G-X.W., C-G.Y., X.Y., K.Z., L.Z.

Planning and Discussion: X-N.Z., X-R.F., Y-S.W., D.C., N.Y., M-J.R., Z.Z., Y.H., X.H., Q.Z., Z-Q.G., L-Z.C., H-M.D., L-Z.C., Q.Z., J-X.Z., H-J.L., M.B., A.C., J.C., X.C., J.D., X.D., Y.D., C.F., T.F., X.F., L-K.G., X.H., C.J., L.L., Q.L., S.L., X.L., J.Q., X-Q.S., G-X.W., H.X., C-G.Y., Z-X.Y., X.Y., K.Z., L.Z.

Implementation and Logistics: X-R.F., N.Y., M-J.R.

Data Collection: X-R.F., N.Y., M-J.R., Z.Z., Y.H., Y-S.W., Q.Z., J-X.Z. Data Informatics: X-R.F., N.Y., Y-S.W., D.C., M-J.R.

Data Analysis: Y-S.W., X-R.F., D.C., H-M.D., Y.H., N.Y., M-J.R., L-Z.C., J-J.N., X.D., B.H., W.H., F.M., Y.W., W.Z.

Initial Drafting of the Manuscript: X-R.F., Y-S.W., D.C. Supervision and Cohort Funding: Q.D., X-N.Z.

Critical Review and Editing of the Manuscript: All authors contributed to the critical review and editing of the manuscript.

## Supplementary Materials

**Figure S1.**
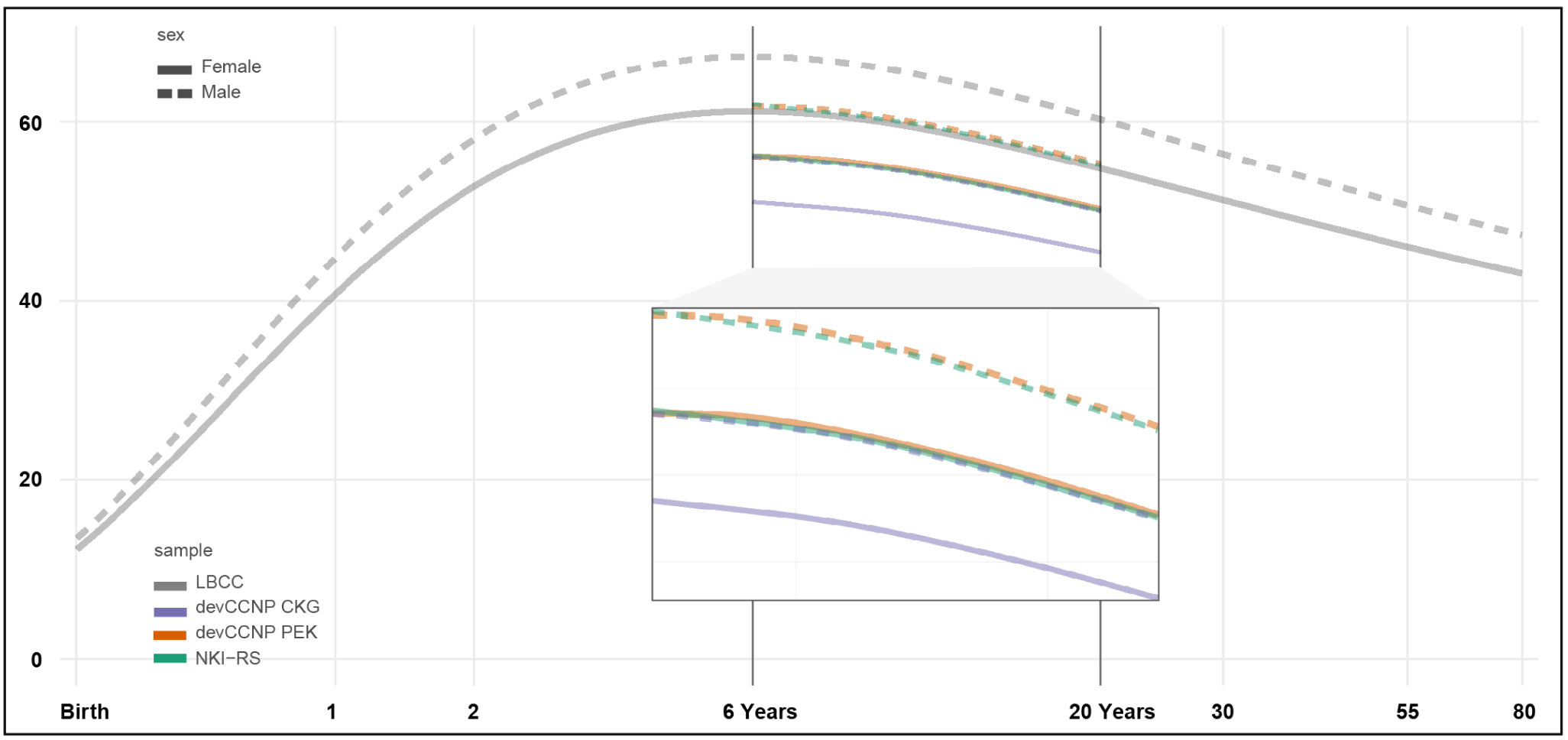
Site/sex-specific brain charts of grey matter volume (GMV). The sex-specific lifespan brain charts of GMV (LBCC, light gray) were adjusted by leveraging the school-aged (6–18 years old) samples for three sites (devCCNP-CKG, purple; devCCNP-PEK, orange; NKI-RS, green), with male (dashed lines) and female (solid lines) respectively. unit: 10*ml* or 10, 000*mm*^3^.

**Figure S2.**
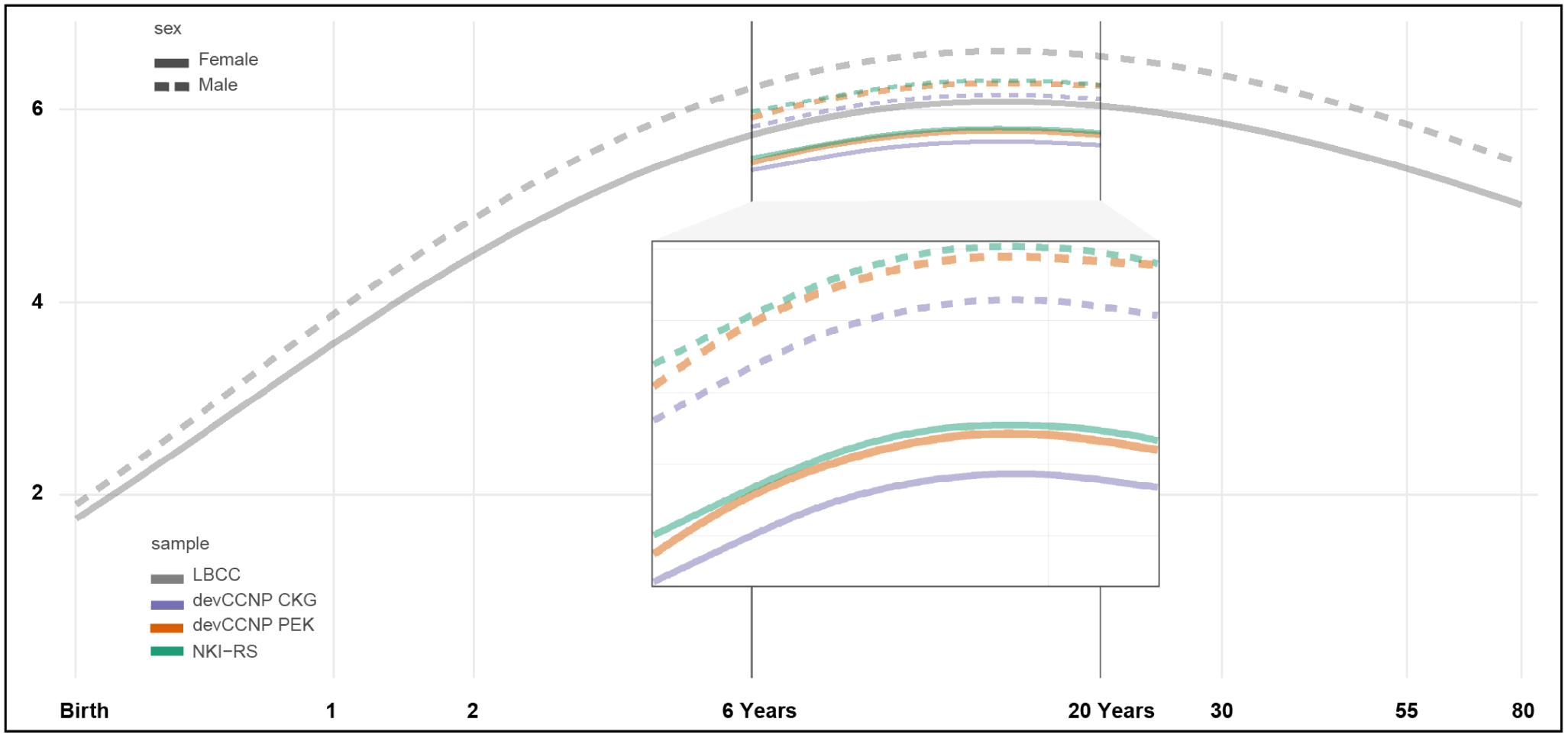
Site/sex-specific brain charts of subcortical grey matter volume (sGMV). The sex-specific lifespan brain charts of sGMV (LBCC, light gray) were adjusted by leveraging the school-aged (6–18 years old) samples for three sites (devCCNP-CKG, purple; devCCNP-PEK, orange; NKI-RS, green), with male (dashed lines) and female (solid lines) respectively. unit: 10*ml* or 10, 000*mm*^3^.

**Figure S3.**
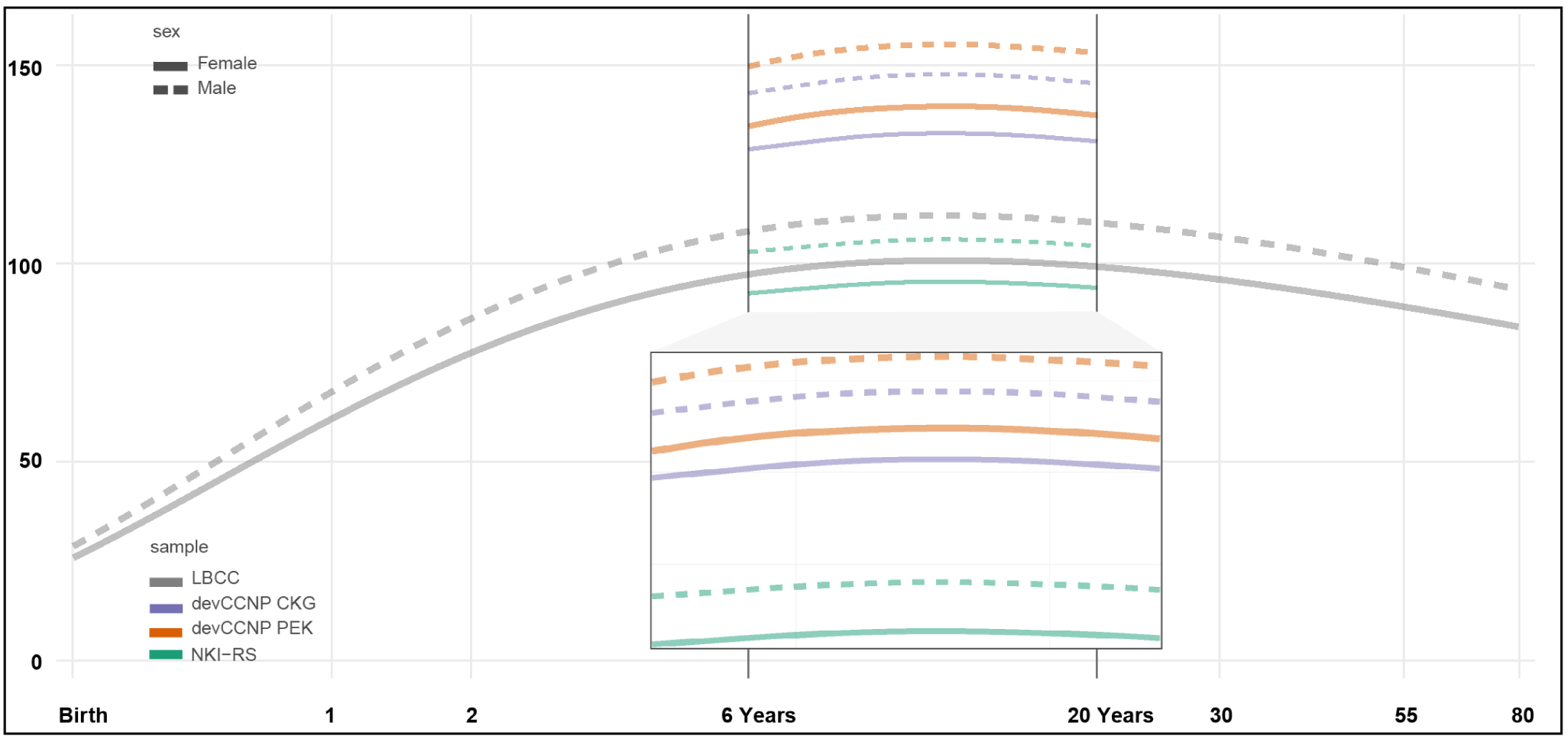
Site/sex-specific brain charts of total cerebrum volume (TCV). The sex-specific lifespan brain charts of TCV (LBCC, light gray) were adjusted by leveraging the school-aged (6–18 years old) samples for three sites (devCCNP-CKG, purple; devCCNP-PEK, orange; NKI-RS, green), with male (dashed lines) and female (solid lines) respectively. unit: 10*ml* or 10, 000*mm*^3^.

**Figure S4.**
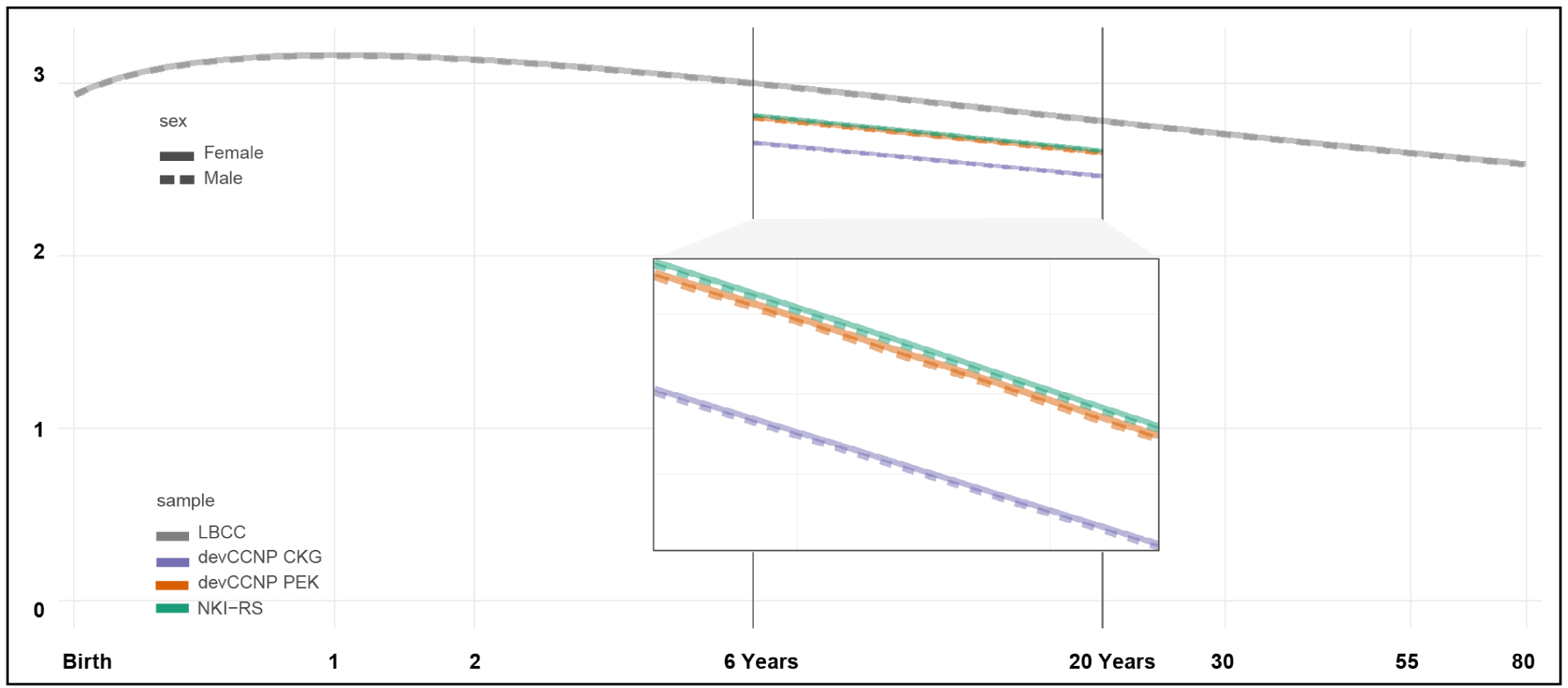
Site/sex-specific brain charts of mean cortical thickness (CT). The sex-specific lifespan brain charts of mean CT (LBCC, light gray) were adjusted by leveraging the school-aged (6–18 years old) samples for three sites (devCCNP-CKG, purple; devCCNP-PEK, orange; NKI-RS, green), with male (dashed lines) and female (solid lines) respectively. unit: *mm*.

**Figure S5.**
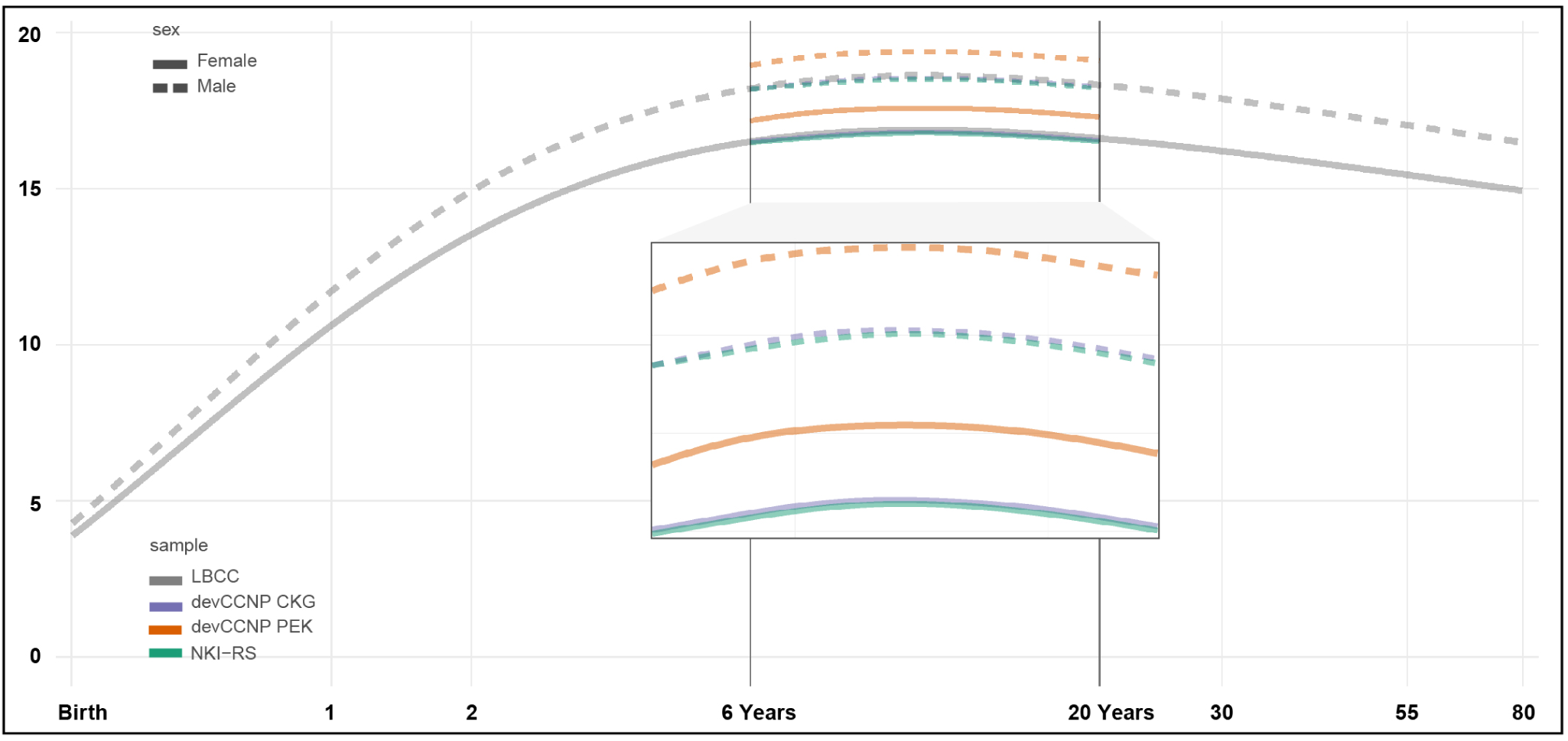
Site/sex-specific brain charts of total surface area (TSA). The sex-specific lifespan brain charts of TSA (LBCC, light gray) were adjusted by leveraging the school-aged (6–18 years old) samples for three sites (devCCNP-CKG, purple; devCCNP-PEK, orange; NKI-RS, green), with male (dashed lines) and female (solid lines) respectively. unit: 10, 000*mm*^2^.

**Figure S6.**
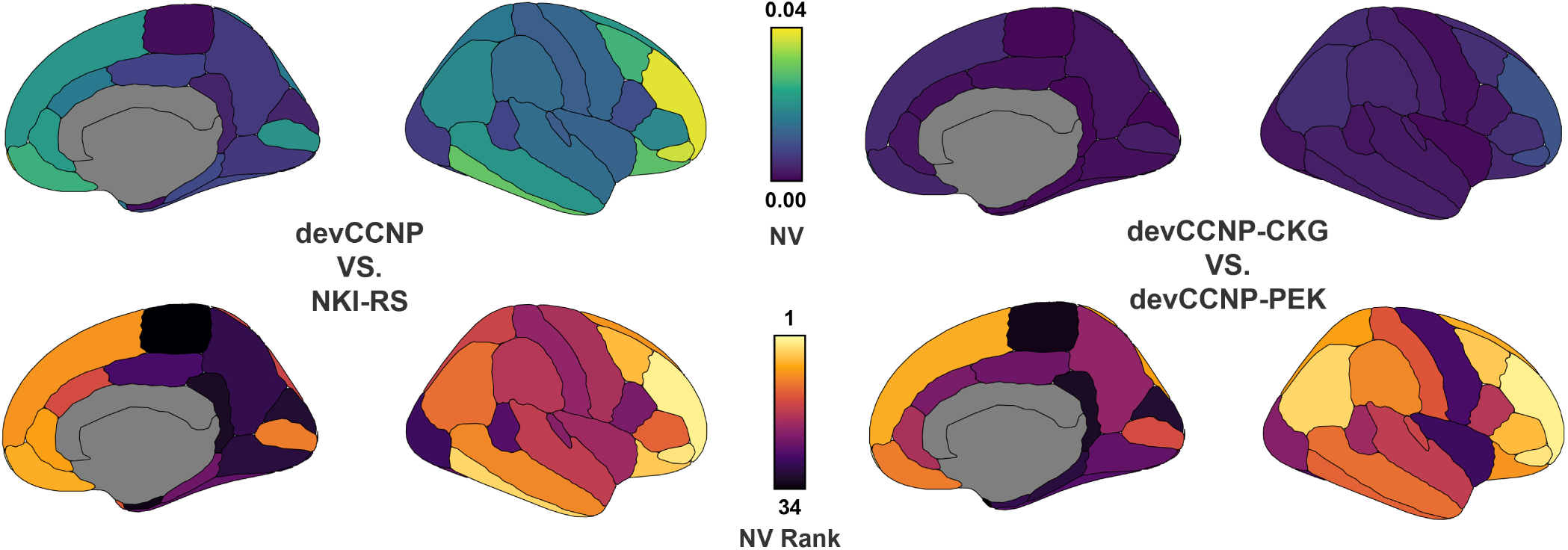
Similarities of brain growth curves between male participants in devCCNP and NKI-RS. NV values of the similarity between the United States and China (top, left) and two Samples within devCCNP (top, right) are presented through 34 gyral-based neuroanatomical regions. NV rank of these parcels are presented respectively bottom) from highest (order 1) to lowest (order 34).

**Table S1.**
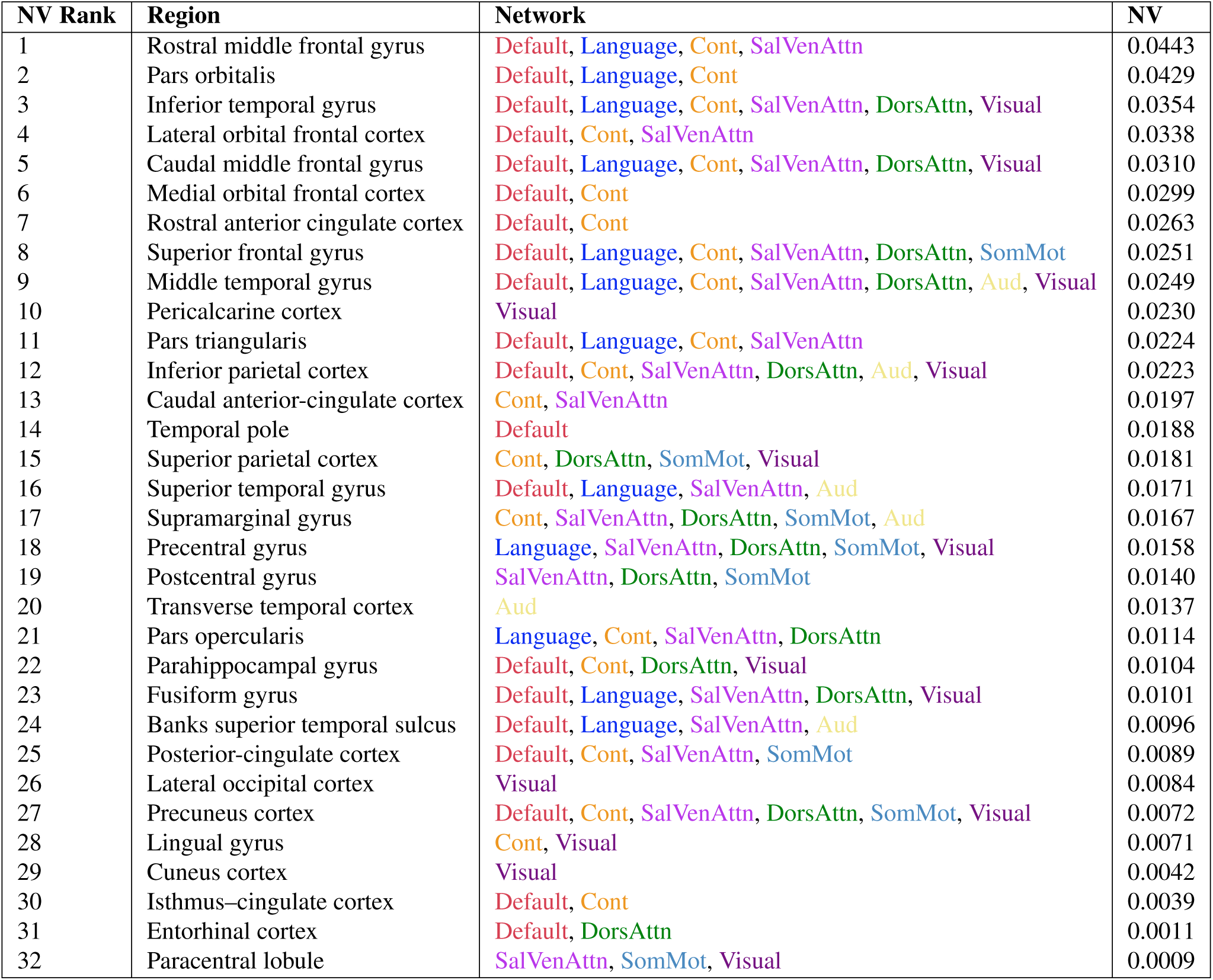
NV among devCCNP and NKI-RS. As explained in the manual delineation procedure of Desikon-Killiany parcellation, the region frontal pole was not actually designed as a measure of the frontal pole itself. Other frontal lobe regions were first designated and the remaining portion was called the frontal pole, which was also proven to be unreliable. The region corpus callosum was introduced to better define the regions around it. Therefore, the NV of these two regions is not shown.

**Table S2.**
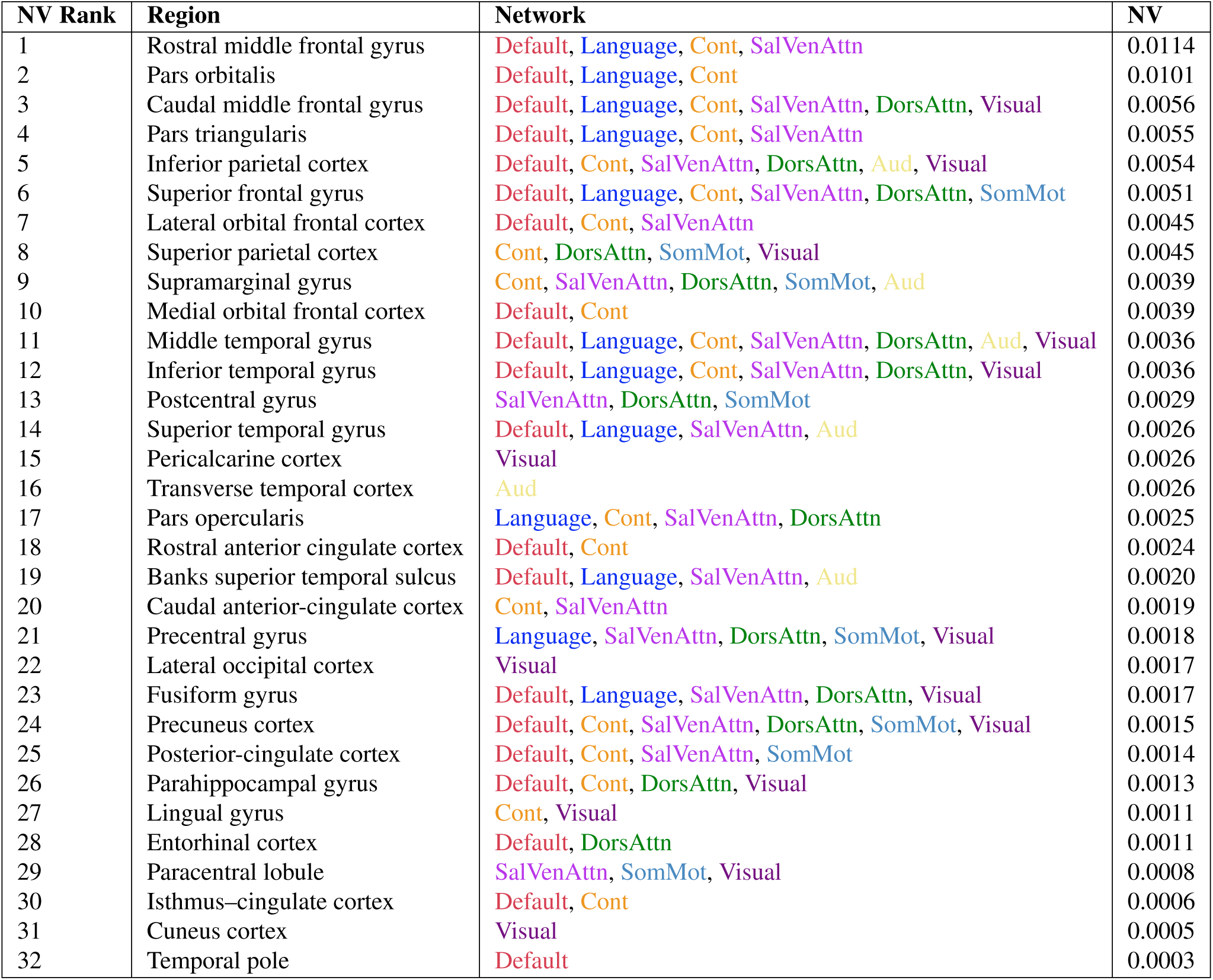
NV among devCCNP-PEK and devCCNP-CKG.

