## Supplementary for "A longitudinal resource for population neuroscience of school-age children and adolescents in China"

### A longitudinal resource for population neuroscience of school-age children and adolescents in China: Supplementary Figures & Tables

**Xue-Ru Fan (范雪如)[1,2,3,†], Yin-Shan Wang (王银山)[1,3,4,†], Da Chang (常达)[1,3,†], Ning Yang (杨宁)[1,2,3,4], Meng-Jie Rong (荣孟杰)[1,2,3,4], Zhe Zhang (张吉吉)[5], Ye He (何叶)[6], Xiaohui Hou (侯晓晖)[7], Quan Zhou (周荃)[1,2,3], Zhu-Qing Gong (宫竹青)[1,2,3], Li-Zhi Cao (曹立智)[2,4], Hao-Ming Dong (董昊铭)[1,4,8,9], Jing-Jing Nie (聂晶晶)[1,3], Li-Zhen Chen (陈丽珍)[1,3], Qing Zhang (张青)[2,4], Jia-Xin Zhang (张家鑫)[2,4], Hui-Jie Li (李会杰)[2,4], Min Bao (鲍敏)[2,4], Antao Chen (陈安涛)[10,11], Jing Chen (陈静)[12,13], Xu Chen (陈旭)[11], Jinfeng Ding (丁金丰)[2,4], Xue Dong (董雪)[2,4], Yi Du (杜忆)[2,4], Chen Feng (冯臣)[2,4], Tingyong Feng (冯廷勇)[11], Xiaolan Fu (傅小兰)[2,14], Li-Kun Ge (盖力锟)[2,4], Bao Hong (洪宝)[12,15], Xiaomeng Hu (胡晓檬)[16], Wenjun Huang (黄文君)[12,15], Chao Jiang (蒋超)[17], Li Li (李黎)[12,13], Qi Li (李琦)[17], Su Li (李甦)[2,4], Xun Liu (刘勋)[2,4], Fan Mo (莫凡)[2,14], Jiang Qiu (邱江)[11], Xue-Quan Su (苏学权)[7], Gao-Xia Wei (魏高峡)[2,4], Yiyang Wu (吴伊扬)[2,4], Haishuo Xia (夏海硕)[11], Chao-Gan Yan (严超赣)[2,4], Zhi-Xiong Yan (颜志雄)[7], Xiaohong Yang (杨晓虹)[16], Wenfang Zhang (张文芳)[2,4], Ke Zhao (赵科)[2,14], Liqi Zhu (朱莉琪)[2,4], Lifespan Brain Chart Consortium (LBCC)[*], Chinese Color Nest Consortium (CCNP)[**], and Xi-Nian Zuo (左西年)[1,2,3,4,7,18,***]**

[1]State Key Laboratory of Cognitive Neuroscience and Learning, Beijing Normal University, Beijing, 100875, China.
[2]Department of Psychology, University of Chinese Academy of Sciences, Beijing, 100049, China.
[3]Developmental Population Neuroscience Research Center, International Data Group/McGovern Institute for Brain Research, Beijing Normal University, Beijing, 100875, China.
[4]Key Laboratory of Behavioural Science, Institute of Psychology, Chinese Academy of Sciences, Beijing, 100101, China.
[5]College of Education, Hebei Normal University, Shijiazhuang, 050024, China.
[6]School of Artificial Intelligence, Beijing University of Posts and Telecommunications, Beijing, 100876, China.
[7]Laboratory of Cognitive Neuroscience and Education, School of Education Science, Nanning Normal University, Nanning, 530299, China.
[8]Changping Laboratory, Beijing, 102206, Chin.a
[9]Department of Psychology, Yale University, New Haven, CT 06511, USA.
[10]School of Psychology, Research Center for Exercise and Brain Science, Shanghai University of Sport, Shanghai, 200438, China.
[11]Faculty of Psychology, Southwest University, Chongqing, 400715, China.
[12]NYU-ECNU Institute of Brain and Cognitive Science at New York University Shanghai, Shanghai, 200062, China.
[13]Faculty of Arts and Science, New York University Shanghai, Shanghai, 200122, China.
[14]State Key Laboratory of Brain and Cognitive Science, Institute of Psychology, Chinese Academy of Sciences, Beijing, 100101, China.
[15]School of Psychology and Cognitive Science, East China Normal University, Shanghai, 200062, China.
[16]Department of Psychology, Renmin University of China, Beijing, 100872, China.
[17]Beijing Key Laboratory of Learning and Cognition, School of Psychology, Capital Normal University, Beijing, 100048, China.
[18]National Basic Science Data Center, Beijing, 100190, China.
[†]These authors contributed equally to this work as first authors
[*]LBCC is an international consortium and has built brain charts to identify previously unreported neurodevelopmental milestones. More information are available at https://github.com/brainchart/lifespan.
[**]CCNP is a long-term effort (2013-2032) to build the lifespan brain-mind development cohort in China, and more consortium information are available at http://deepneuro.bnu.edu.cn/?p=163.

#### Supplementary Materials

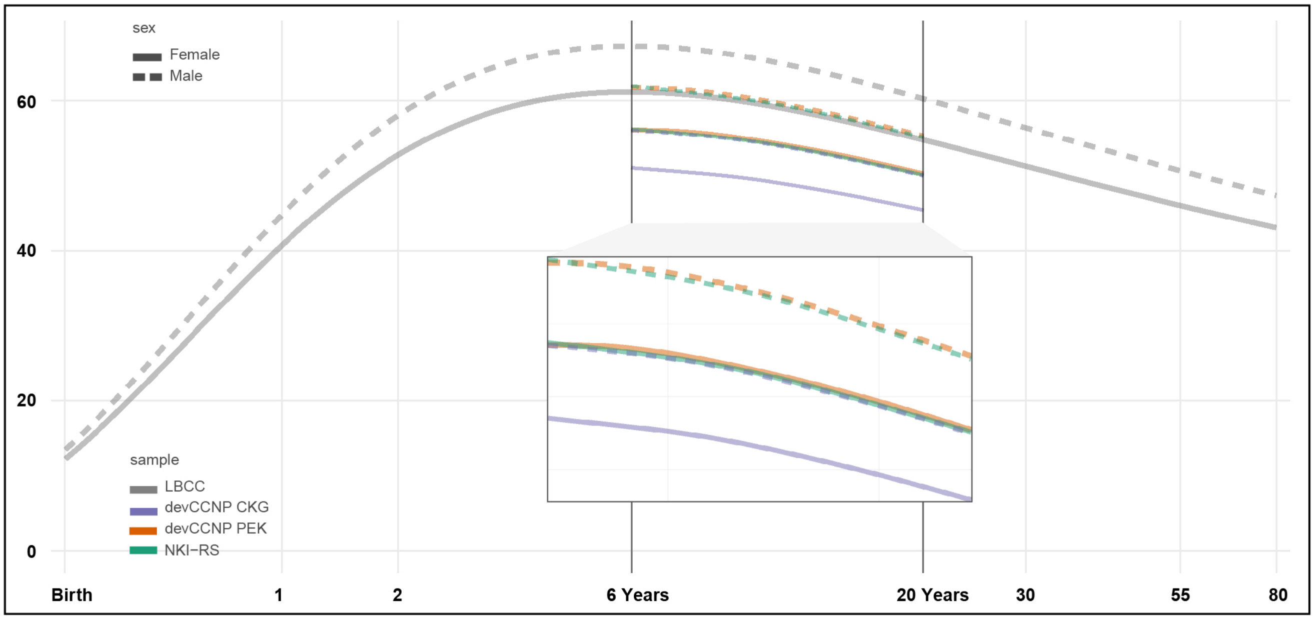


**Figure S1. Site/sex-specific brain charts of grey matter volume (GMV)**. The sex-specific lifespan brain charts of GMV (LBCC, light gray) were adjusted by leveraging the school-aged (6–18 years old) samples for three sites (devCCNP-CKG, purple; devCCNP-PEK, orange; NKI-RS, green), with male (dashed lines) and female (solid lines) respectively. unit: $10ml$ or $10,000mm^3$.

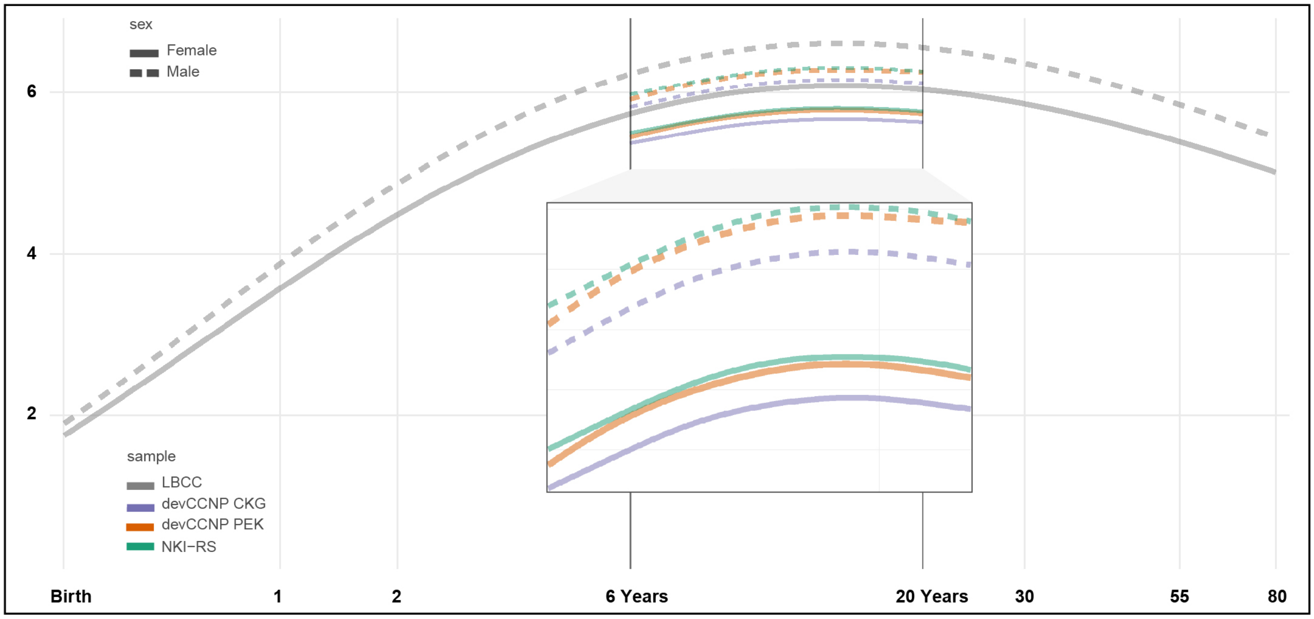


**Figure S2. Site/sex-specific brain charts of subcortical grey matter volume (sGMV)**. The sex-specific lifespan brain charts of sGMV (LBCC, light gray) were adjusted by leveraging the school-aged (6–18 years old) samples for three sites (devCCNP-CKG, purple; devCCNP-PEK, orange; NKI-RS, green), with male (dashed lines) and female (solid lines) respectively. unit: $10ml$ or $10,000mm^3$.

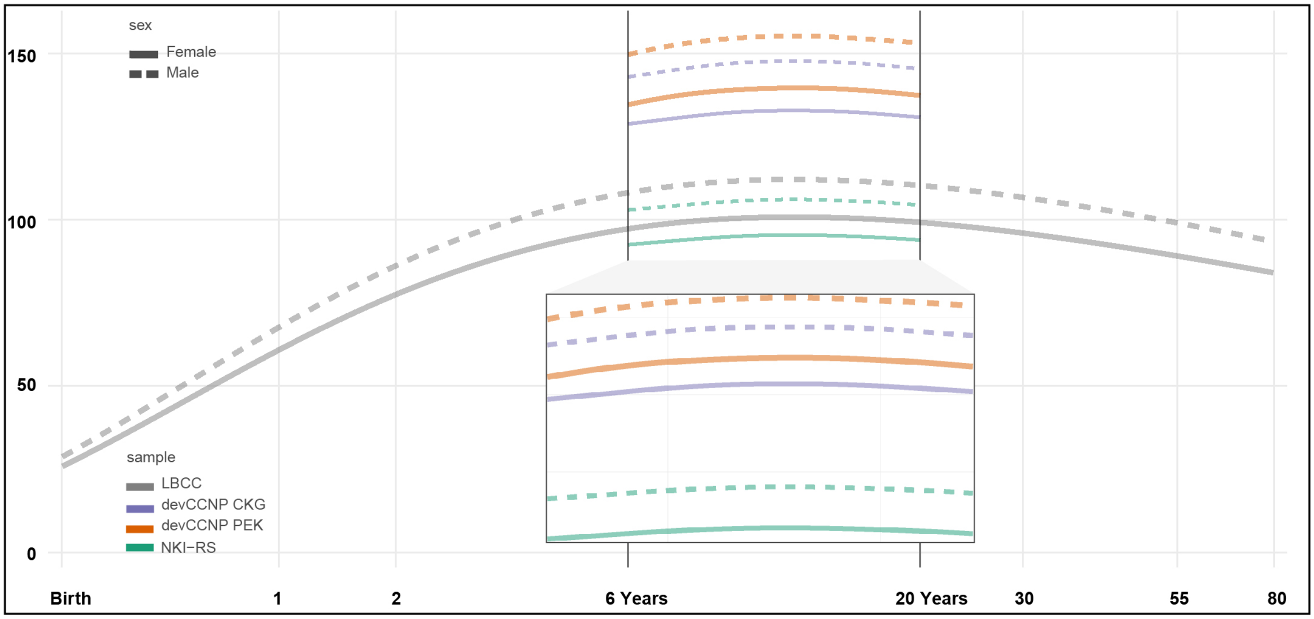


**Figure S3.** **Site/sex-specific brain charts of total cerebrum volume (TCV)**. The sex-specific lifespan brain charts of TCV (LBCC, light gray) were adjusted by leveraging the school-aged (6–18 years old) samples for three sites (devCCNP-CKG, purple; devCCNP-PEK, orange; NKI-RS, green), with male (dashed lines) and female (solid lines) respectively. unit: $10ml$ or $10,000mm^3$.

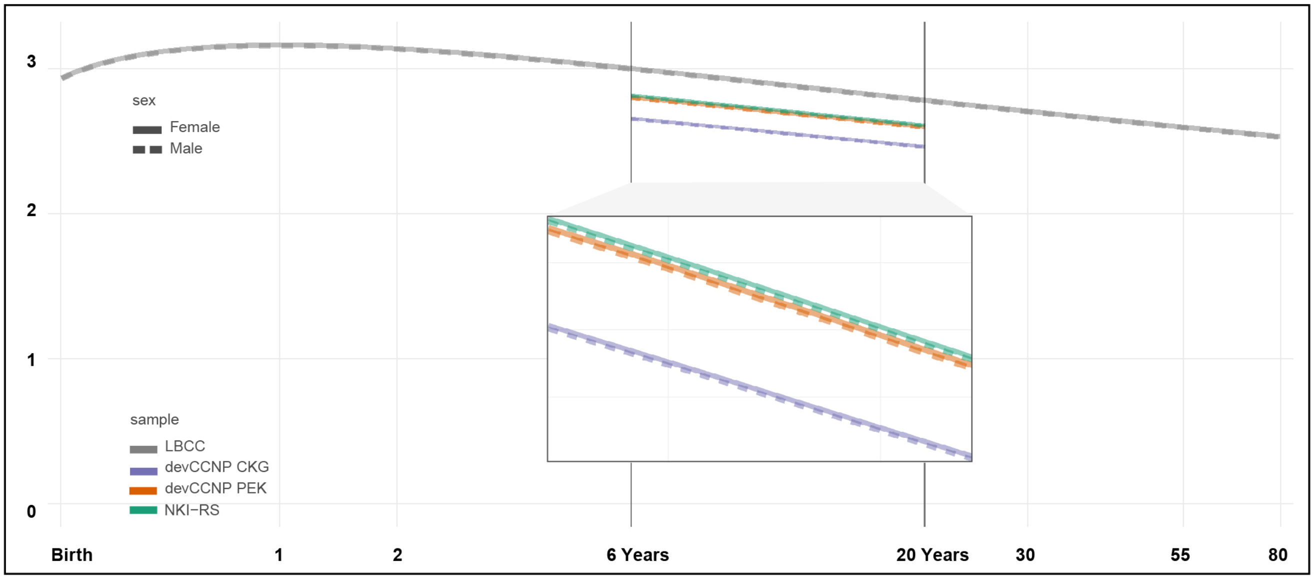


**Figure S4.** **Site/sex-specific brain charts of mean cortical thickness (CT)**. The sex-specific lifespan brain charts of mean CT (LBCC, light gray) were adjusted by leveraging the school-aged (6–18 years old) samples for three sites (devCCNP-CKG, purple; devCCNP-PEK, orange; NKI-RS, green), with male (dashed lines) and female (solid lines) respectively. unit: *mm*.

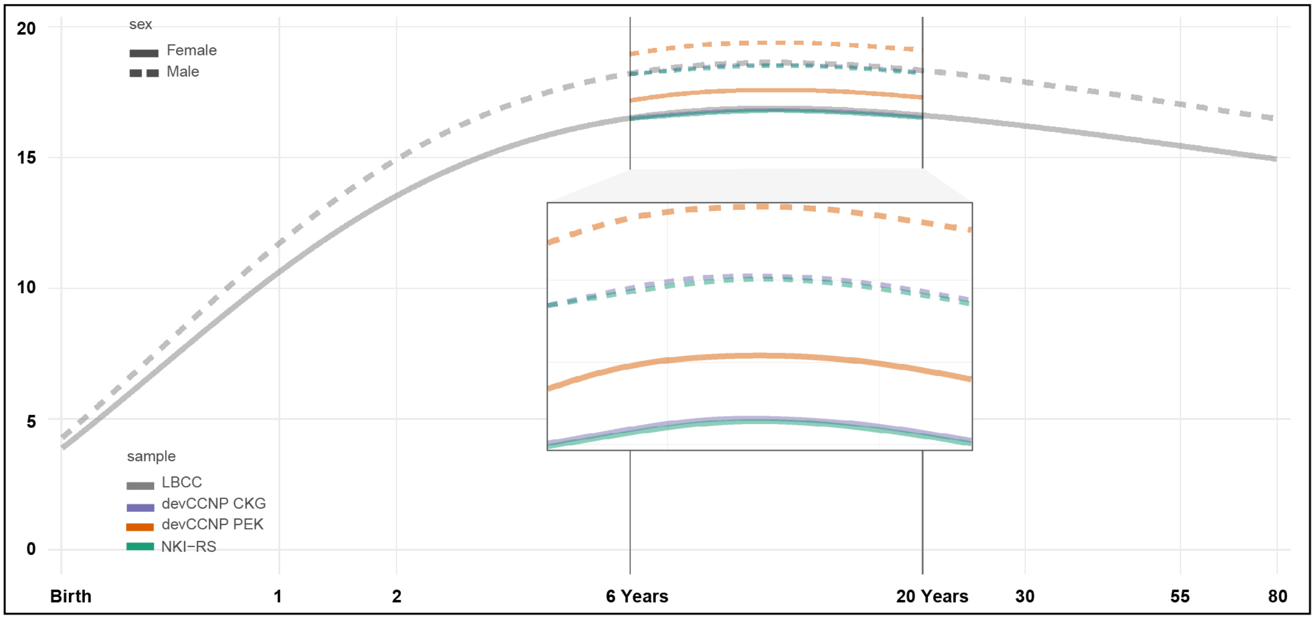


**Figure S5. Site/sex-specific brain charts of total surface area (TSA).** The sex-specific lifespan brain charts of TSA (LBCC, light gray) were adjusted by leveraging the school-aged (6–18 years old) samples for three sites (devCCNP-CKG, purple; devCCNP-PEK, orange; NKI-RS, green), with male (dashed lines) and female (solid lines) respectively. unit: $10,000mm^2$.

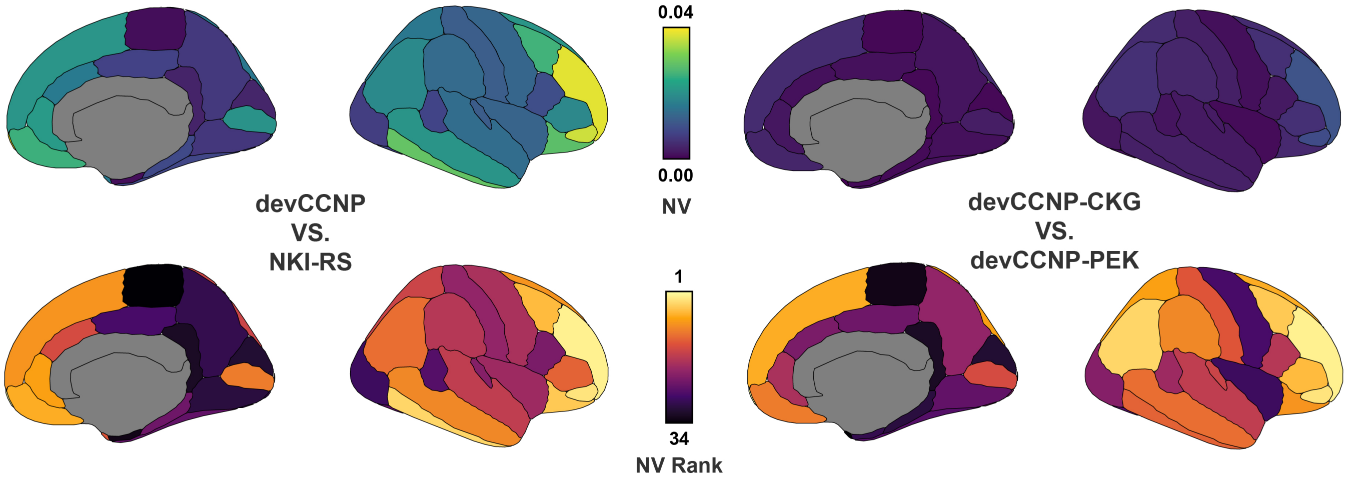


**Figure S6. Similarities of brain growth curves between male participants in devCCNP and NKI-RS.** NV values of the similarity between the United States and China (top, left) and two Samples within devCCNP (top, right) are presented through 34 gyral-based neuroanatomical regions. NV rank of these parcels are presented respectively bottom) from highest (order 1) to lowest (order 34).

| NV Rank | Region | Network | NV |
|---|---|---|---|
| 1 | Rostral middle frontal gyrus | Default, Language, Cont, SalVenAttn | 0.0443 |
| 2 | Pars orbitalis | Default, Language, Cont | 0.0429 |
| 3 | Inferior temporal gyrus | Default, Language, Cont, SalVenAttn, DorsAttn, Visual | 0.0354 |
| 4 | Lateral orbital frontal cortex | Default, Cont, SalVenAttn | 0.0338 |
| 5 | Caudal middle frontal gyrus | Default, Language, Cont, SalVenAttn, DorsAttn, Visual | 0.0310 |
| 6 | Medial orbital frontal cortex | Default, Cont | 0.0299 |
| 7 | Rostral anterior cingulate cortex | Default, Cont | 0.0263 |
| 8 | Superior frontal gyrus | Default, Language, Cont, SalVenAttn, DorsAttn, SomMot | 0.0251 |
| 9 | Middle temporal gyrus | Default, Language, Cont, SalVenAttn, DorsAttn, Aud, Visual | 0.0249 |
| 10 | Pericalcarine cortex | Visual | 0.0230 |
| 11 | Pars triangularis | Default, Language, Cont, SalVenAttn | 0.0224 |
| 12 | Inferior parietal cortex | Default, Cont, SalVenAttn, DorsAttn, Aud, Visual | 0.0223 |
| 13 | Caudal anterior-cingulate cortex | Cont, SalVenAttn | 0.0197 |
| 14 | Temporal pole | Default | 0.0188 |
| 15 | Superior parietal cortex | Cont, DorsAttn, SomMot, Visual | 0.0181 |
| 16 | Superior temporal gyrus | Default, Language, SalVenAttn, Aud | 0.0171 |
| 17 | Supramarginal gyrus | Cont, SalVenAttn, DorsAttn, SomMot, Aud | 0.0167 |
| 18 | Precentral gyrus | Language, SalVenAttn, DorsAttn, SomMot, Visual | 0.0158 |
| 19 | Postcentral gyrus | SalVenAttn, DorsAttn, SomMot | 0.0140 |
| 20 | Transverse temporal cortex | Aud | 0.0137 |
| 21 | Pars opercularis | Language, Cont, SalVenAttn, DorsAttn | 0.0114 |
| 22 | Parahippocampal gyrus | Default, Cont, DorsAttn, Visual | 0.0104 |
| 23 | Fusiform gyrus | Default, Language, SalVenAttn, DorsAttn, Visual | 0.0101 |
| 24 | Banks superior temporal sulcus | Default, Language, SalVenAttn, Aud | 0.0096 |
| 25 | Posterior-cingulate cortex | Default, Cont, SalVenAttn, SomMot | 0.0089 |
| 26 | Lateral occipital cortex | Visual | 0.0084 |
| 27 | Precuneus cortex | Default, Cont, SalVenAttn, DorsAttn, SomMot, Visual | 0.0072 |
| 28 | Lingual gyrus | Cont, Visual | 0.0071 |
| 29 | Cuneus cortex | Visual | 0.0042 |
| 30 | Isthmus–cingulate cortex | Default, Cont | 0.0039 |
| 31 | Entorhinal cortex | Default, DorsAttn | 0.0011 |
| 32 | Paracentral lobule | SalVenAttn, SomMot, Visual | 0.0009 |

**Table S1. NV among devCCNP and NKI-RS**. As explained in the manual delineation procedure of Desikon-Killiany parcellation, the region frontal pole was not actually designed as a measure of the frontal pole itself. Other frontal lobe regions were first designated and the remaining portion was called the frontal pole, which was also proven to be unreliable. The region corpus callosum was introduced to better define the regions around it. Therefore, the NV of these two regions is not shown.

| NV Rank | Region | Network | NV |
|---|---|---|---|
| 1 | Rostral middle frontal gyrus | Default, Language, Cont, SalVenAttn | 0.0114 |
| 2 | Pars orbitalis | Default, Language, Cont | 0.0101 |
| 3 | Caudal middle frontal gyrus | Default, Language, Cont, SalVenAttn, DorsAttn, Visual | 0.0056 |
| 4 | Pars triangularis | Default, Language, Cont, SalVenAttn | 0.0055 |
| 5 | Inferior parietal cortex | Default, Cont, SalVenAttn, DorsAttn, Aud, Visual | 0.0054 |
| 6 | Superior frontal gyrus | Default, Language, Cont, SalVenAttn, DorsAttn, SomMot | 0.0051 |
| 7 | Lateral orbital frontal cortex | Default, Cont, SalVenAttn | 0.0045 |
| 8 | Superior parietal cortex | Cont, DorsAttn, SomMot, Visual | 0.0045 |
| 9 | Supramarginal gyrus | Cont, SalVenAttn, DorsAttn, SomMot, Aud | 0.0039 |
| 10 | Medial orbital frontal cortex | Default, Cont | 0.0039 |
| 11 | Middle temporal gyrus | Default, Language, Cont, SalVenAttn, DorsAttn, Aud, Visual | 0.0036 |
| 12 | Inferior temporal gyrus | Default, Language, Cont, SalVenAttn, DorsAttn, Visual | 0.0036 |
| 13 | Postcentral gyrus | SalVenAttn, DorsAttn, SomMot | 0.0029 |
| 14 | Superior temporal gyrus | Default, Language, SalVenAttn, Aud | 0.0026 |
| 15 | Pericalcarine cortex | Visual | 0.0026 |
| 16 | Transverse temporal cortex | Aud | 0.0026 |
| 17 | Pars opercularis | Language, Cont, SalVenAttn, DorsAttn | 0.0025 |
| 18 | Rostral anterior cingulate cortex | Default, Cont | 0.0024 |
| 19 | Banks superior temporal sulcus | Default, Language, SalVenAttn, Aud | 0.0020 |
| 20 | Caudal anterior-cingulate cortex | Cont, SalVenAttn | 0.0019 |
| 21 | Precentral gyrus | Language, SalVenAttn, DorsAttn, SomMot, Visual | 0.0018 |
| 22 | Lateral occipital cortex | Visual | 0.0017 |
| 23 | Fusiform gyrus | Default, Language, SalVenAttn, DorsAttn, Visual | 0.0017 |
| 24 | Precuneus cortex | Default, Cont, SalVenAttn, DorsAttn, SomMot, Visual | 0.0015 |
| 25 | Posterior-cingulate cortex | Default, Cont, SalVenAttn, SomMot | 0.0014 |
| 26 | Parahippocampal gyrus | Default, Cont, DorsAttn, Visual | 0.0013 |
| 27 | Lingual gyrus | Cont, Visual | 0.0011 |
| 28 | Entorhinal cortex | Default, DorsAttn | 0.0011 |
| 29 | Paracentral lobule | SalVenAttn, SomMot, Visual | 0.0008 |
| 30 | Isthmus–cingulate cortex | Default, Cont | 0.0006 |
| 31 | Cuneus cortex | Visual | 0.0005 |
| 32 | Temporal pole | Default | 0.0003 |

**Table S2.** NV among devCCNP-PEK and devCCNP-CKG
